# Prioritization of abiotic and biotic plant stress responses through ABI1 phosphatase and CPK5 calcium-dependent protein kinase switch

**DOI:** 10.1101/839662

**Authors:** Heike Seybold, Jennifer Bortlik, Xiyuan Jiang, Anja Liese, Benjamin Conrads, Wolfgang Hoehenwarter, Susanne Matschi, Tina Romeis

**Author notes:** Department of Plant and Environmental Sciences, Alexander Silberman Institute of Life Sciences, The Hebrew University of Jerusalem, Givat Ram, 9190401 Jerusalem, Israel (Heike Seybold). Leibniz Institute of Vegetable and Ornamental Crops, 14979 Großbeeren, Germany (Jennifer Bortlik).

## Abstract

In nature plants are constantly challenged by simultaneous abiotic and biotic stresses, and under conflicting stress scenarios prioritization of stress responses is required for plant survival. Calcium-dependent protein kinase CPK5 is a central hub in local and distal immune signaling, required for hormone salicylic acid (SA)-dependent immunity and pathogen resistance. Here we show that CPK5-dependent immune responses and pathogen resistance are inhibited upon abscisic acid (ABA) treatment or in genetic mutant backgrounds lacking PP2C phosphatase activities including *abi1-2*, whereas immune responses are enhanced by co-expression of active ABI1 phosphatase variants. Biochemical studies and mass spectrometry-based phospho-site analysis reveal a direct ABI1 phosphatase-catalyzed de-phosphorylation of CPK5 auto-phosphorylation site T98. Mimicking continuous de-phosphorylation in CPK5_T98A_ leads to enhanced ROS production and more resistant plants, mimicking the auto-phosphorylated status in CPK5_T98D_, reduces CPK5-mediated immune responses. Mechanistic insight identifies differential phosphorylation at T98 in the N-terminal domain of CPK5 to control the level of interaction between the kinase and its substrate protein rather than CPK5 catalytic activity. Thus, CPK5-catalyzed immune signaling may become discontinued even at an elevated cytoplasmic calcium concentration.

Our work reveals an elegant mechanism for stress response prioritization in plants: The ABA-dependent phosphatase ABI1, negative regulator of abiotic responses, functions as positive regulator of biotic stress responses, stabilizing CPK5-dependent immune signaling in the absence of ABA. Continuous pathogen survey activates plant immunity in environmentally friendly conditions, whereas under severe abiotic stress the phosphatase/kinase pair prohibits immune signaling through a direct biochemical switch involving two key regulatory enzymes of these antagonistic pathways.

**Significance Statement:** Plants challenged by simultaneous abiotic and biotic stresses must prioritize in conflicting scenarios to guarantee survival. Pathogen resistance and immune memory depends on the phytohormone salicylic acid (SA). Adaptation to abiotic stress signaling involves the phytohormone abscisic acid (ABA). We identify a direct biochemical switch by which ABA-mediated abiotic signaling prioritizes over SA-dependent immune responses via reversible phosphorylation at a single protein mark involving two key regulatory enzymes of these antagonistic pathways. Phosphatase ABI1 de-phosphorylates calcium-dependent protein kinase CPK5 at an auto-phosphorylation site T98, which effects the interaction efficiency between the kinase and its substrate. Under abiotic stress ABA mediates phosphatase inhibition, which facilitates prolonged auto-phosphorylation of CPK5, preventing CPK5 substrate interaction and ultimately stop CPK5-mediated immune signaling.

## Introduction

Plants experience a continuously changing environment and must permanently adapt to new conditions. These encompass abiotic changes like temperature variations, shortage in water supply, or high salinity, and biotic challenges, where plants have to tackle attacks by various microbial pathogens or herbivorous insects. Because stress responses are energy consuming and often come to the cost of a retardation or a full stop of growth (1, 2), it is in the interest of a plant to implement mechanisms to limit specific stress responses in time and to prioritize between multiple simultaneous external challenges.

Pathogen infections lead to a rapid activation of the plant immune system. Plants perceive attacking microbes in a non-species-specific manner via molecules, Pathogen-Associated Molecular Patterns (PAMPs), which bind as ligands to corresponding receptors (Pattern-Recognition Receptors, PRRs) on the cell surface, intracellular defense reactions become initiated and lead to PAMP-triggered immunity (PTI). These cytoplasmic reactions include an increase in the concentration of calcium ions (Ca^2+^), the activation of protein kinases including Ca^2+^-decoding calcium-dependent protein kinases (CDPKs), production of reactive oxygen species (ROS), phytohormone signaling, transcriptional reprogramming, and changes in metabolites to ultimately accomplish pathogen resistance (3). A second layer of defense has evolved in plants, Effector-Triggered Immunity (ETI), which detects PTI-suppressing effectors from adapted pathogens. Intracellular recognition inducing ETI results in the induction of a similar set of reactions compared to PTI, except immune responses are stronger and long-lasting and lead to a hypersensitive cell death. Remarkably, ETI has recently been associated with a sudden and a longer-lasting increase in the cytoplasmic Ca^2+^ concentration mediated by cation permeable pores (4–7). Additionally, PTI and ETI may prime a systemic long-term immune memory, which prepares the plant for consecutive pathogen attacks (8–10). Calcium-dependent protein kinase CPK5 from Arabidopsis is a highly Ca^2+^-sensitive CDPK expressed in plant mesophyll cells and functions as a key regulatory enzyme in plant immunity (11–15). CPK5 contributes to local pathogen resistance, to defense signal spread, and to the onset and maintenance of long-term resistance. Plants displaying enhanced CPK5-signaling develop stronger and prolonged resistance in PTI and ETI (12, 16), and respective plants accumulate high levels of SA, display stunted growth and develop lesions indicative for a constitutively activated immune system (12, 16). Consistently, plants which code for non-functional *CPK5* protein variants lack prolonged immunity in an ETI-mimicking autoimmune phenotype (17). In a biological context, CPK5 immune signaling may be stopped by a direct biochemical inactivation of the enzyme. CDPKs are modular enzymes, in which a protein kinase effector domain is directly linked via a pseudo-substrate region to a calcium-binding sensor domain, the so-called calcium activation domain (CAD), which in CPK5 contains four consensus EF-hand Ca^2+^-binding motifs (18–20). Ca^2+^ binding induces a conformational change into an open enzyme conformation (step 1), which is prerequisite for ATP-dependent trans-phosphorylation (step 2) (12, 18, 21, 22). For CPK5 the activation threshold has been determined to Kd50 [Ca^2+^] of 100 nM, an intracellular Ca^2+^ concentration close to a plant cell’s resting state (16). This low Ca^2+^ value implies that a gradual withdrawal of Ca^2+^ may be insufficient for a rapid biochemical inactivation of the catalytic CPK5 activity, in particular under conditions of ETI. Thus, additional control mechanisms are required for rapid enzyme inactivation, for example through post-translational modifications and/or via a control of substrate protein access.

Abscisic acid (ABA) is the major phytohormone to trigger abiotic stress tolerance, causal to the induction of rapid stomatal closure (23, 24). The control of ABA function depends on the biosynthesis of the phytohormone, on the presence of ABA receptor proteins, and on downstream signaling components. Key regulators of the ABA signal transduction pathways involve clade 2 phosphatases (PP2Cs) such as ABI1 or its close homologues HAB1, or PP2CA, and protein kinases such as OST1 plus other (Ca^2+^-regulated) protein kinases (25, 26). In the absence of an abiotic challenge to the plant, phosphatase ABI1 functions as a negative regulator of ABA signaling. Drought and increased ABA levels lead to an ABA/ABA-receptor-mediated sequestration and inactivation of the PP2C phosphatase, to release phosphorylated active OST1, and to further progress ABA signal transduction (27–29). ABA signaling induces ion channel-mediated stomatal closure (30–32). ABA-dependent closure of guard cells has also been discussed as pre-invasive immunity, an important layer in PTI against invading *Pto* DC3000 (33–36). Stomata close within one hour in response to PAMPs and a parallel integration of SA- and ABA-signaling was observed (37, 38). However, prolonged stomatal closure prohibits gas exchange and compromises key functions in guard cell physiology with respect to photosynthesis and transpiration (32). Thus, long-term and systemic pathogen resistance of post-invasive immunity in mesophyll cells may employ an antagonistic relation of SA- and ABA-signaling.

When exposed simultaneously to biotic pathogen- and abiotic water shortage stress, drought may represent the higher threat for the survival of an entire plant than a pathogen attack which may be limited to defined plant parts. Thus, ABA suppresses plant defense responses although a pathogen has invaded plant tissue. Exogenous application of ABA increased the susceptibility of plants to bacterial as well as fungal pathogens (39–45). Also, ABA-deficient or -insensitive mutants were more resistant towards (hemi-) biotrophic pathogens (42–44, 46). In these trade-offs between biotic and abiotic stress responses the plant prioritizes the abiotic stress responses (47). Prioritization mechanisms against immunity may occur as early as at the level of perception, for example through a control of the secretion and accessibility of PRR receptors in the plasma membrane (48). At the signaling level, ABA-induced transcriptional activation of phosphatase genes, with respective phosphatases inactivating immune-associated MAP kinase pathways (49). At late stages, prioritization can be implemented by employing a phytohormone-mediated cross-talk, in which ABA signaling suppresses SA synthesis/signaling and blocking SA-dependent immunity (50, 51).

Here we decipher a novel direct biochemical mechanism involving reversible phosphorylation at a protein mark, which controls the switch between biotic to abiotic stress responses. ABI1-catalyzed de-phosphorylation of an auto-phosphorylation site of CPK5 links the two antagonistic stress response pathways. CPK5, master regulator for biotic SA-dependent immune signaling is linked to the phosphatase ABI1, a negative regulator of abiotic ABA-signaling, and in the absence of severe abiotic stress a positive regulator of immune signaling.

## Results

### ABA reverts CPK5 signaling-dependent immune phenotype

Enhanced CPK5 signaling in CPK5-YFP #7 plants increases local immune responses and bacterial resistance in young plants and results in SA-dependent long-term immunity (9, 12, 16, 17). Furthermore, in later plant stages enhanced CPK5 signaling correlates with a reduced plant rosette size and a SA-dependent lesion mimic phenotype (Fig. S1A-C). Significant size differences became evident from day 33 and further increased with plant age (Fig. S1A). Interestingly, continuous treatment with 10 μM ABA over several weeks with ABA application every second day reverted all CPK5 signaling-dependent growth phenotypes of CPK5-YFP #7, and plants were of wild type size with no or only few lesions (Fig. 1A-D, Fig. S1C). Transcript levels of the SA biosynthesis and signaling marker genes *ICS1* and *PR1*, and of SA-independent CPK5 marker gene *NHL10* were completely reverted (Fig. 1 E-G). In CPK5-YFP #7, highly elevated SA levels (about 8-fold in 6 and 10 weeks-old plants) is not mirrored by basal endogenous ABA phytohormone levels (1.5 – 2-fold induction) suggesting that these endogenous ABA levels are not directly dependent on SA (Fig. S2).

**Fig. 1.**
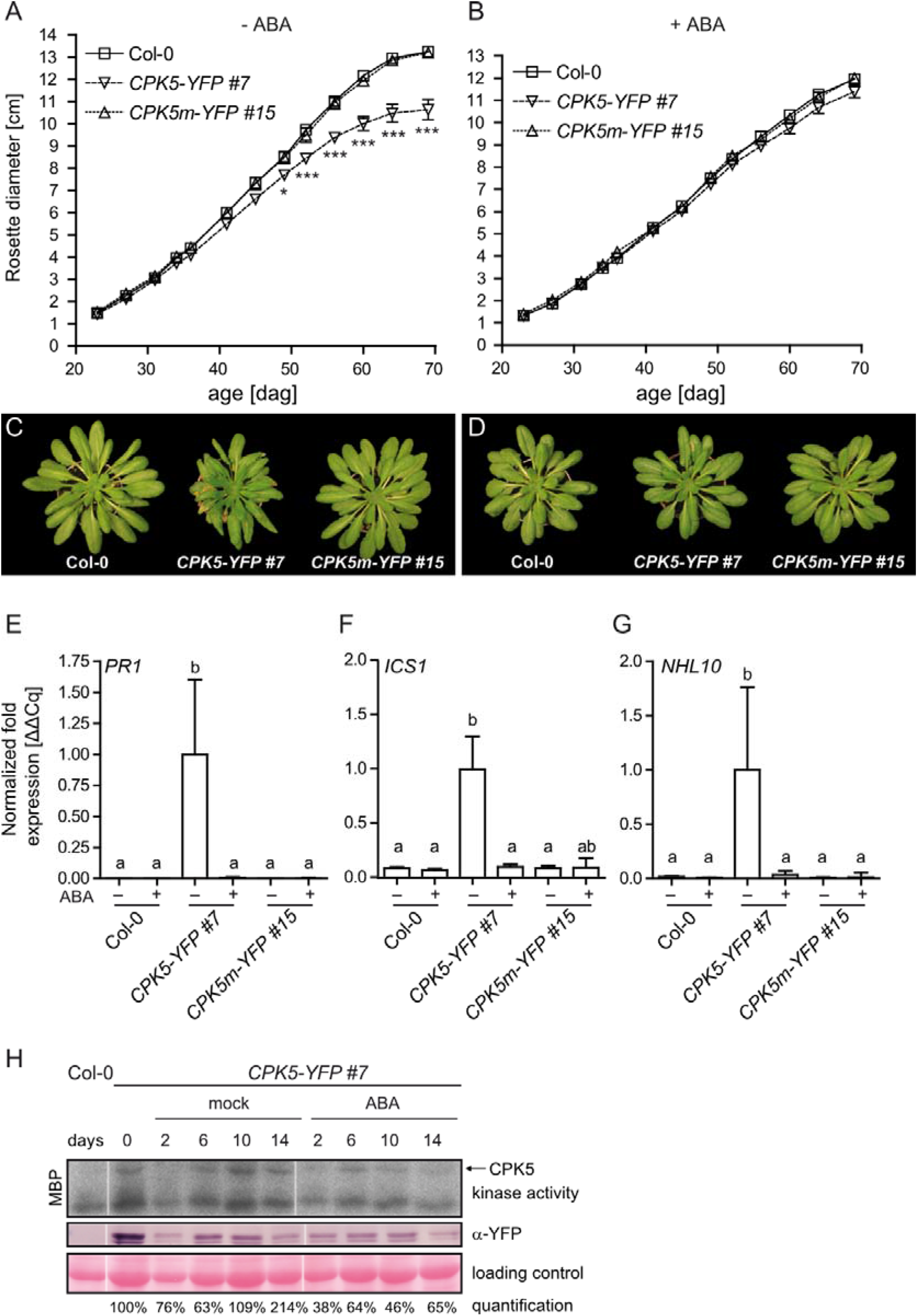
ABA inhibits the CPK5-YFP phenotype and reduces kinase phosphorylation. (*A-D*) Monitoring of rosette diameter during growth phase of Col-0, *CPK5-YFP #7* and *CPK5m-YFP #15 (kinase-deficient variant*) (n = 8-13) over a time course of 10 weeks and corresponding phenotypes of 10 weeks old plants in the absence (*A, C*) and presence (*B, D*) of treatment with 10 μM ABA. (*E-G*) Basal transcript levels of marker genes in 6 weeks old plants as shown in (*A*) with or without ABA treatment (n =4). (*H*) *In-gel* kinase using MPB of protein extracts originating from overexpression line *CPK5-YFP #7* after spray treatment with 3 μM ABA. Kinase phosphorylation activity was assessed over a period of 14 days with repeated ABA treatment every second day. Expression of CPK5 was detected via western blot analysis. Enzyme transphosphorylation of MBP was quantified by ImageJ analysis and activity levels were normalized to the control sample at 0 days without treatment, after normalization of transphosphorylation band to YFP protein levels of the respective sample. Bars represent mean ± SD of n biological replicates. Data in *A, B, and E-G* were evaluated for statistical differences using two-way ANOVA (Bonferroni post-test against all groups). Significant differences * = p < 0.05, ** = p < 0.01 and *** = p < 0.001. Significantly different groups were assigned different letters.

Biochemical assessment of plant crude extracts in an *in-gel* kinase assay (Fig. 1H) reveals that ABA treatment between 2 - 14 days of plants does neither significantly alter the CPK5 protein amount, when comparing the western blot signal (middle panel) to the (unregular) RuBisCO loading control (lower panel), nor its migration in a lower mobility band indicative for phosphorylation at the CPK5 protein itself (middle panel) (Fig. 1H, middle and lower panel). However, when quantifying and normalizing the MBP (trans-)phosphorylation signal (upper panel) to the YFP signal as indicated by the numbers below, an overall reduced phosphorylation strength is observed for example at days 10 and 14 (above 100 % and 200 % in mock treatment; above 40 and 60 % upon ABA treatment).

### ABI1 regulates CPK5 function and phosphorylation activity

To assess whether ABA signaling-associated PP2C-type of protein phosphatases are mechanistically involved in the loss of CPK5 trans-phosphorylation signal, we generated crosses between CPK5-YFP #7 and mutant lines *abi1-2*, *hab1-1*, and *pp2ca*, lacking respective protein phosphatase activities. CPK5-YFP protein expression in the resulting lines has been confirmed by western blot and by detection of eYFP transcripts (Fig. S3 B,C). In the absence of these phosphatase activities, elevated transcript levels of *PR1* and *NHL10* and the lesion mimic phenotype, both hallmark for enhanced CPK5 immune signaling, was reverted in all lines (Fig. 2 A,B and Fig. S3A), and despite slightly less CPK5-YFP protein present in the cross to *abi1-2* (Fig. S3B). In bacterial growth assays, the CPK5-dependent constitutive resistance to bacterial pathogens *Pto* DC3000 in CPK5-YFP #7 was entirely reverted or at least partially lost to wild type when CPK5-YFP is expressed in the genetic backgrounds of the phosphatase mutants (Fig. 2C). Consistently, a reduction of rapid flg22-induced ROS generation occurs in the phosphatase mutant background reminiscent to *cpk5* mutant plants (Fig. S3D) (12). These data suggest that phosphatase activity is beneficial to CPK5 function.

**Fig. 2.**
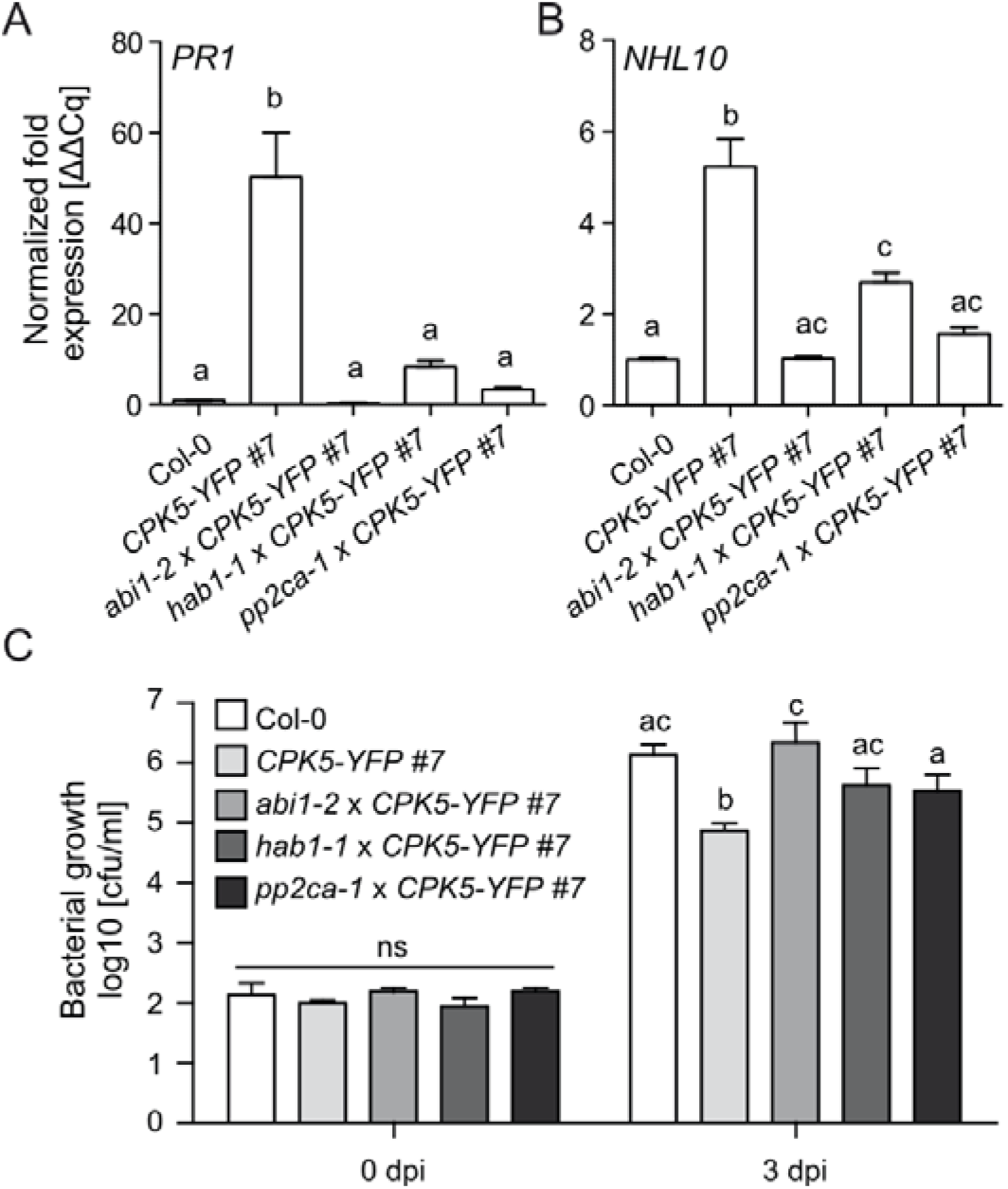
CPK5-dependent resistance phenotypes are reverted in *PP2C* knock out backgrounds. (*A, B*) Basal transcript levels of defense related marker genes *PR1 (A*) and *NHL10 (B*) in Col-0, overexpression line *CPK5-YFP #7* and crossings of this line to *PP2C* knockout mutants *abi1-2*, *hab1-1* and *pp2ca-1* (bars represent mean ± SE, n = 5-11). (*C*) Bacterial growth of *Pto* DC3000 at 0 and 3 days after infection in overexpression lines as in (*A, B*) (bars represent mean ± SE, n = 8-14). Data were analyzed for statistical differences using one-way ANOVA (*A, B*) or two-way ANOVA (*C*) with Bonferroni post-tests comparing all groups, p < 0.05. Significantly different groups were assigned different letters, ns = no significant differences.

To address this positive effect directly, we transiently co-expressed active ABI1 phosphatase variants and CPK5 kinase together with NADPH oxidase RBOHD in *Nicotiana (N.) benthamiana* and compared the levels of CPK5-dependent RBOHD-catalyzed ROS production in the absence of the phosphatase (12). We used the truncated constitutive active variant CPK5-VK (12), lacking its regulatory calcium activation domain (CAD), which is a biochemically active enzyme independent of further stimulation. Constitutive phosphatase activity mimicking the absence of ABA was achieved using two gain-of-function ABA insensitive phosphatase variants derived from the semi-dominant *abi1-1* and *abi1-11* alleles. ABI1-1 carries the amino acid substitution G180D (52, 53) and ABI1-11 the substitution G181S (54). The presence of both phosphatase variants, ABI1-1 and ABI1-11, together with CPK5-VK induced a dramatic increase in ROS production (Fig. 3A), which is still evident in the absence of ectopic RBOHD, relying on endogenous RBOH enzymes. No ROS production occurs when these phosphatase variants ABI1-1 and ABI1-11 are combined with RBOHD alone, indicating that the positive ABI phosphatase effect requires CPK5.

**Fig. 3.**
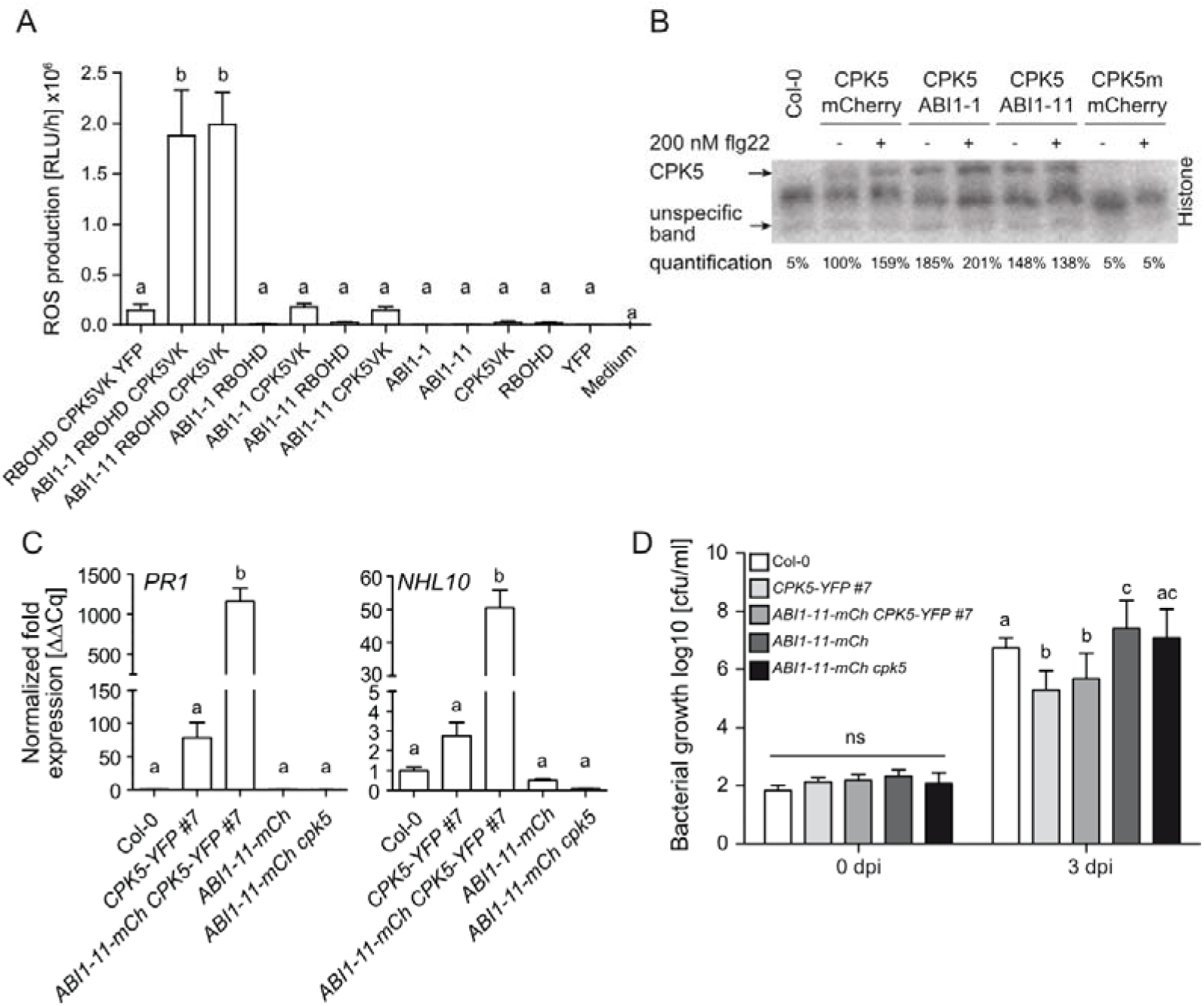
ABI1 promotes CPK5-dependent immune responses and kinase phosphorylation. (*A*) Constitutively active versions ABI1-1-mCherry and ABI1-11-mCherry were transiently co-expressed with different combinations of RBOHD-YFP and CPK5-VK-StrepII in *N. benthamiana* (n = 8). ROS were measured with a luminol-based assay, values represent accumulated ROS over 1 hour. (*B*) *In-gel* kinase assay using histone of protein extracts from Arabidopsis mesophyll protoplast transiently co-expressing CPK5-StrepII with ABI1-1-mCherry, ABI1-11-mCherry or mCherry-tag alone as well as kinase-deficient CPK5m-Strep as negative control. Protoplasts were treated with 200 nM flg22 or a control treatment for 10 min. Enzyme transphosphorylation of histone was quantified by ImageJ analysis and activity levels were normalized to the indicated unspecific band. (*C*) Basal transcript levels of defense related marker genes *PR1* and *NHL10* in Col-0 and the overexpression lines *CPK5-YFP #7, ABI1-11-mCherry CPK5-YFP, ABI1-11-mcherry* and *ABI1-11-mCherry cpk5* (n = 4). (*D*) Bacterial growth of *Pto* DC3000 tested directly and 3 days after infection in overexpression lines as mentioned in (*C*) (n =14). Values represent mean ± SD of n biological replicates. Data were evaluated for statistical differences using one-way ANOVA (*A, C*) or two-way ANOVA (*D*) with Bonferroni post-tests comparing all groups, p < 0.05. Significantly different groups were assigned different letters, ns = no significant differences.

To biochemically assess CPK5 (trans-) phosphorylation activity we conducted an *in-gel* kinase assay with crude extracts originating from Arabidopsis mesophyll protoplasts transiently expressing and co-expressing full-length CPK5 plus ABI1-1 or ABI1-11. In the absence of further stimulation, the quantified phosphorylation signal (normalized to an unspecific band) set to 100 % for CPK5 increased to 185 % for ABI1-1 and 148 % for ABI1-11, whereas no signal is detectable in kinase-deficient CPK5m extracts. Treatment with flg22 leads to a further increase to 159 % for CPK5 and 201 % for the stronger ABI1-1 allele (Fig. 3B). These data suggest a more efficient phosphorylation of the histone protein by CPK5 in the presence of active ABI1 phosphatase, with ABI1-1 displaying a stronger effect than ABI1-11.

### ABI1 positively influences CPK5-dependent immunity

To further corroborate an ABI1/CPK5 relation, we generated stable transgenic lines which co-express phosphatase mCherry fusion proteins of ABI1-1-mCh and ABI1-11-mCh together with CPK5-YFP by dipping the ABI1 constructs into the CPK5-YFP #7 background. As appropriate controls, we transformed Col-0 and *cpk5* mutant with these ABI1 constructs. Co-expression of ABI1-1 with CPK5-YFP resulted in severely dwarfed seedlings, which rarely survived. Plant lines co-expressing ABI1-11-mCh together with CPK5-YFP were viable. The expression of proteins ABI1-11-mCh and CPK5-YFP was verified by western blot in all lines (Fig. S4A) and by quantitative real-time PCR (qPCR) (Fig. S4B). Co-expression of ABI1-11-mCh with CPK5-YFP resulted in a more severe lesion mimic phenotype (Fig. S4C) accompanied by strongly increased *PR1* and *NHL10* transcript levels (Fig. 3C), even further enhanced in dwarfed *ABI1-1-mCh x CPK5-YFP* plants (Fig. S5A). In bacterial growth assays both CPK5-YFP #7 and *ABI1-11-mCh x CPK5-YFP* were equally resistant to *Pto* DC3000 (Fig. 3D). In contrast, plants expressing ABI1-11-mCherry in *cpk5* or Col-0 background showed wild type-like susceptibility.

### ABI1 catalyzes direct de-phosphorylation at phosphorylation site T98

We next assessed whether CPK5 is a functional direct substrate for ABI1 for de-phosphorylation. Recombinant tagged CPK5 protein was subjected to autocatalysis. Kinase activity was then inhibited by K252-a, and the reaction was divided and subsequently further incubated over a time course in the presence of either active or heat-inactivated ABI1 phosphatase. Active phosphatase reduced the CPK5 auto-phosphorylation signal to 23 % within 60 minutes (Fig. S6A). A considerably weaker decrease in band intensity occurred with heat-inactivated ABI1. These data suggest that ABI1 catalyzes the de-phosphorylation of CPK5 at one/some autocatalytic phosphorylation sites but likely not all. To identify amino acids targeted by ABI1-catalyzed de-phosphorylation in CPK5, the experiment was repeated with CPK5-YFP followed by mass spectrometry-based differential phospho-site analysis. Because ABI1 triggered an increase in ROS production together with the CPK5-VK variant (Fig. 3A) we focused on the N-terminal part of the CPK5 protein encompassing the variable and the kinase domain. From the four detected phosphorylated amino acids at S8, T98, S100 and S338 only T98 revealed a differential phosphorylation pattern with a decrease in intensity in the presence of active phosphatase (Fig. 4A, Fig. S6 B, C). To validate T98 as a regulatory phospho-site for CPK5 function recombinant CPK5 protein was incubated with either active ABI1 or a phosphatase-inactive variant ABI1-D177A. MS-based targeted phospho-site analysis via parallel reaction monitoring (PRM) revealed a site-specific de-phosphorylation of pT98 by ABI1 but not of pS100 (Fig. 4B, Fig. S6D).

**Fig. 4.**
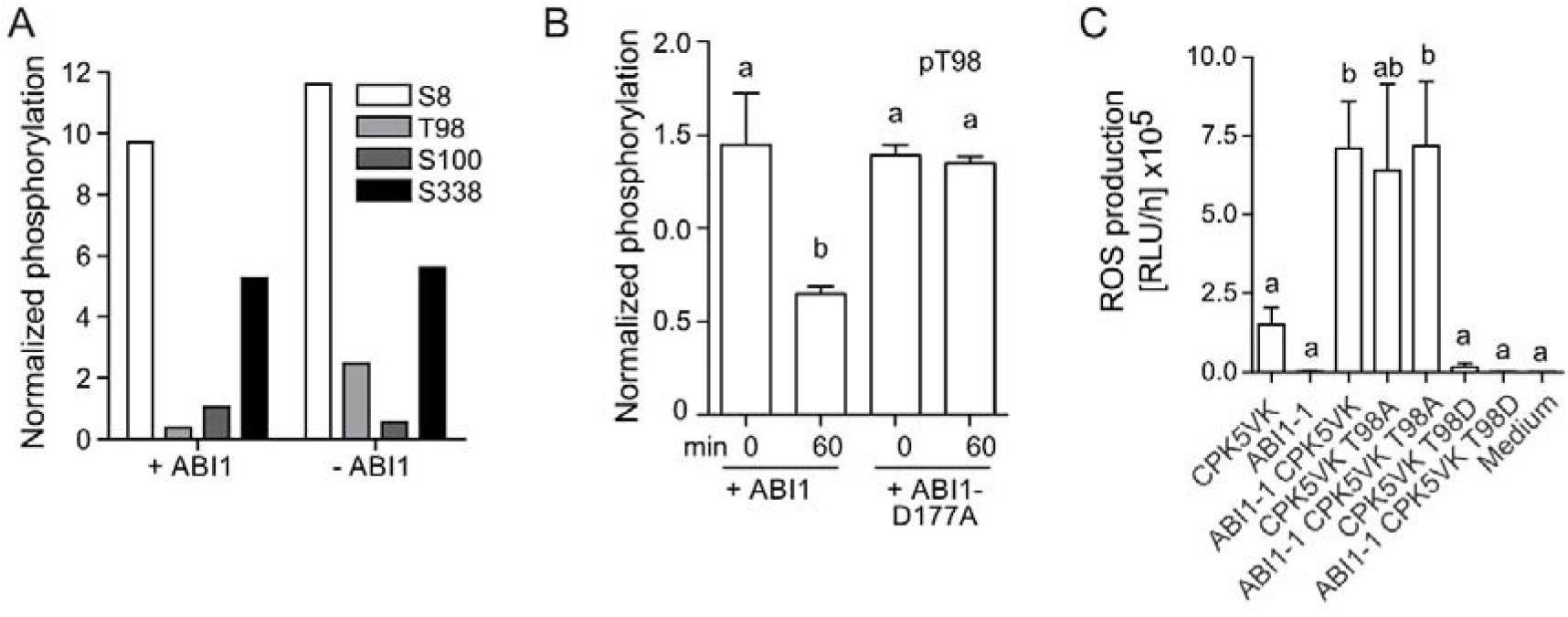
CPK5 auto-phosphorylation site T98 is targeted by ABI1 and is required for CPK5 function in ROS production. (*A*) Quantitative MS-screen of *recombinant auto-phosphorylated* CPK5-YFP in the absence or presence of catalytically active ABI1. MS measurements results from two experiments each of *At*CPK5 with and without ABI1 were concatenated. In each of the concatenated results, the ratio of the number of pT98 peptide spectral matches (PSM) to the number of PSM of non-phosphorylated counterpart peptides was used to quantify site specific phosphorylation, and normalized to the fraction of total *At*CPK5 #PSM of total recorded MS2 spectra in each of the concatenated results. (*B*) Targeted MS-analysis of T98 phosphorylation upon ABI1-treatment. Recombinant auto-phosphorylated CPK5-StrepII was incubated with active ABI1 or catalytically inactive ABI1-D177A for 0 or 60 min. For quantification of phosphorylation, the summed area under the curve (AUC) of fragment ions was used and relative abundance of phosphorylated peptides compared to all peptides spanning the modification site were calculated. (*C*) Transient ROS assay as shown in Fig. 3A with phospho-abolishing mutant (T98A) and phospho-mimic (T98D) versions of CPK5-VK (n = 8). Values represent mean ± SE of n biological replicates. Data were evaluated for statistical differences using one-way ANOVA with Bonferroni post-tests comparing all groups, p < 0.05. Significantly different groups were assigned different letters.

### CPK5 phosphorylation site T98 mediates immune signaling

To further investigate the role of T98 we generated CPK5 variants carrying the amino acid substitutions T98A (mimicking constant de-phosphorylation through ABI1), and T98D (mimicking constant CPK5 auto-phosphorylation in the absence of (active) ABI1), and enzyme activity was assessed in a transient ROS assay in *N. benthamiana* as shown in Fig. 3A. Remarkably, a dramatic increase in transient ROS production is observed with CPK5 T98A alone, in the absence of the phosphatase, to the level of CPK5-VK (wild type sequence) and ABI1-1 co-expression, and no further increase was obtained in the presence of ABI1-1 (Fig. 4C). In contrast, no ROS production occurs with the CPK5 T98D variant, also not in the presence of the ABI1-1 phosphatase. These data confirm T98 as a crucial amino acid in CPK5, which controls CPK5 function in immune signaling.

We next generated transgenic Arabidopsis lines which express the wild type CPK5-YFP, and the variants CPK5_T98A_-YFP or CPK5_T98D_-YFP as fusion proteins in the *cpk5* mutant background and *CPK5* transcript expression was confirmed (Fig. 5B, Fig. S7A, B). Lines of about equally low CPK5 protein levels were selected by western blot to allow a comparison of the T98 substitution. This is of particular importance because we have demonstrated that the protein amount of active CPK5 enzyme may result in enhanced CPK5 signaling influencing the degree of defense responses from as early as Ca^2+^ signature (55) to defense gene expression and resistance to bacterial pathogen *Pto* DC 3000 (12). A smaller rosette size in 6 weeks-old plants was observed upon expression of CPK5-YFP compared to the *cpk5* parental line, which is even further reduced in the T98A variant but not in the T98D variant (Fig. 5A). Consistently to the rosette sizes, expression of defense marker gene *NHL10* is elevated in T98A lines, compared to CPK5 wild type, T98D or to *cpk5* (Fig. S7C). In these newly generated lines with an overall low (~ 10 %) CPK5-YFP protein expression level compared to *CPK5-YFP #7*, both T98A lines are more resistant to *Pto* DC3000 in bacterial growth assays than the parental *cpk5* line or lines expressing the CPK5 wild type or the T98D variant (Fig. 5C). These data corroborate T98 as a crucial amino acid in CPK5, which directs function in immune signaling and resistance, suitable for a mechanistic switch between biotic vs. abiotic stress responses.

**Fig. 5.**
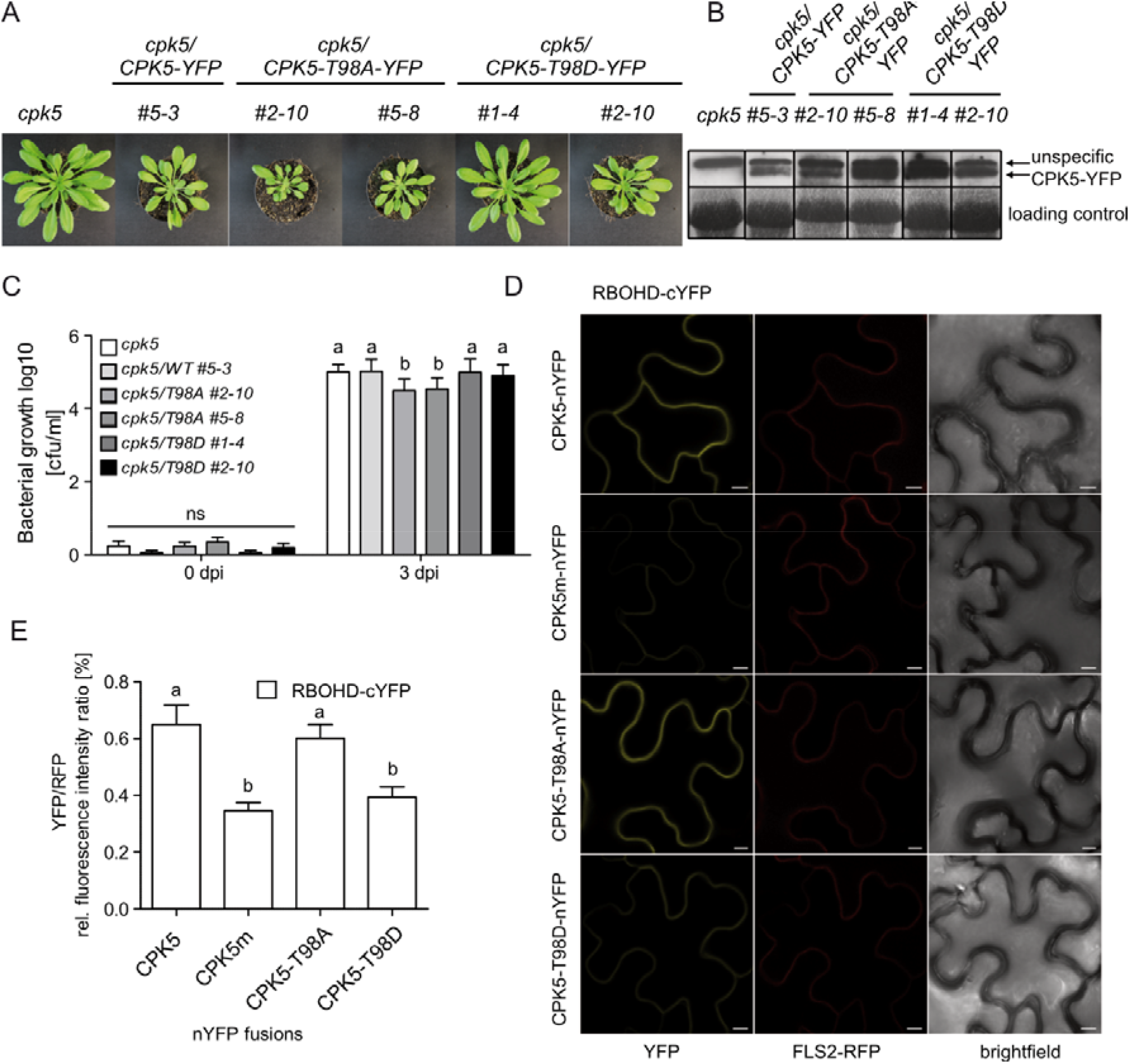
T98 phosphorylation site is essential for in planta-function of CPK5. (*A*) Phenotype of plant rosettes of 6 weeks old plants and (*B*) confirmed protein expression by western blot of CPK5 wild-type, phospho-mutant (T98A) or phospho-mimic (T98D) variants in a *cpk5* mutant background. (*C*) Bacterial growth of *Pto* DC3000 tested directly and 3 days after infection in overexpression lines as in (*A*) (n = 8-12). (*D, E*) Evaluation of *in-vivo* interaction of CPK5 and RBOHD by bimolecular fluorescence complementation assay (BiFC) in *N. benthamiana. (E*) Quantification of reconstituted YFP fluorescence signals: YFP and RFP channels were quantified via ImageJ, and RFP fluorescence signal was used as normalization factor (n=20). Values represent mean ± SD of n biological replicates. Data were analyzed for statistical differences using two-way ANOVA (*C*) or one-way ANOVA (*E*) with Bonferroni post-tests comparing all groups, p < 0.05. Significantly different groups were assigned different letters, ns = no significant differences. Scale bar = 20 μM.

### The CPK5 phosphorylation site T98 mediates kinase-substrate interaction

To gain mechanistic insight in the role of T98 for CPK5 function we compared kinase activity in *in-vitro* assays using short peptide substrates (16). No difference neither in calcium-dependency nor in phosphorylation activity was observed with recombinant purified CPK5 protein, irrespectively of the T98A or the T98D substitution toward 10 amino acid RBOHD peptide nor to Syntide2 (Fig. S8).

We next analyzed protein interaction between CPK5 and full-length RBOHD protein, an *in-vivo* substrate protein of CPK5, using bimolecular fluorescence complementation (BiFC). Different variants of CPK5-nYFP were transiently co-expressed with RBOHD-cYFP in *N. benthamiana* and protein amounts were confirmed (Fig. 5D; Fig. S9A). Both CPK5 wild type and the T98A variants were able to interact with RBOHD (Fig. 5D), yet significantly reduced interaction occurred with the T98D or when kinase-deficient CPK5m was assessed (Fig. 5E). Results were additionally confirmed via an independent BiFC experimental setup, in which RBOHD and CPK5 variants were expressed from a single binary plasmid (Fig. S9B-D (56). The obtained RBOHD/CPK5 protein-protein interaction data are consistent with those showing CPK5-and RBOHD-mediated ROS production (Figs. 3A, 4C). These data indicate that CPK5 T98 site influences kinase trans-phosphorylation efficiency and subsequent immune responses through access to the phosphorylation substrate rather than a direct control of enzyme catalytic activity.

Taken together these data suggest that the auto-phosphorylation site T98 of CPK5 is perceptive to ABI1-catalyzed de-phosphorylation. The phosphorylation status of T98 controls CPK5 function. We present (de-)phosphorylation of T98 as a mechanistic switch between biotic and abiotic stress signaling pathways.

## Discussion

Plant survival depends on stress response activation and inactivation. Under simultaneous multiple stress challenges, a plant prioritizes between different abiotic and biotic danger scenarios. Here we have identified a pair of key regulatory proteins: a calcium sensor protein kinase, CPK5, positive regulator of immune signaling, and protein phosphatase, ABI1, negative regulator of ABA- and abiotic stress signaling. Both enzymes are mechanistically linked via reversible phosphorylation at CPK5 T98 to orchestrate stress signaling and resource allocation.

CPK5 becomes biochemically activated within sec to min upon pathogen perception and functions as signaling hub in the control of the plant immune response. In local immunity enhanced CPK5 signaling in CPK5-YFP overexpressing lines leads to an enforcement of the cytoplasmic Ca^2+^ signature (55), ROS generation, defense gene expression and pathogen resistance (12). Furthermore, CPK5 contributes to post-invasive immunity during the onset and maintenance of systemic acquired resistance in the entire plant (12, 16, 19). These features make CPK5 a most suitable target for the plant to switch into and out of immune signaling when surveying and prioritizing between biotic and abiotic danger scenarios. CPK5 protein is a highly Ca^2+^ responsive enzyme with a low threshold for the Ca^2+^ binding-induced conformational change prerequisite to enzyme activation. This low threshold for activity may also be the reason for the enzyme’s multiple functional contribution to defense signal initiation, propagation and resistance. However, under conditions of severe abiotic challenge the plant may have to stop (CPK5-mediated) immune reactions. Biochemical inactivation of CPK5 through withdrawal of Ca^2+^ may neither be fast nor efficient, particularly not, when concomitant abiotic stress independently elevates cytoplasmic calcium levels.

The reversible post-translational (auto-)phosphorylation at CPK5 T98 represents an elegant intrinsic biochemical mechanism to stop CPK5 function and interfere with immune signaling.

We could demonstrate that the phosphorylated form displays less efficient interaction between the CPK5 kinase and its protein substrate. Thus, despite of potentially elevated cytoplasmic Ca^2+^ levels which promote a Ca^2+^-bound open CPK5 conformation and catalytic activity, CPK5-catalyzed trans-phosphorylation and subsequent downstream signal transduction is discontinued.

Under environmentally non-challenging conditions, ongoing immune survey and signaling correlates with and depends on a constant de-phosphorylation of CPK5 T98, catalyzed by PP2C phosphatases, as in the context of ABA signaling by ABI1 (Fig. 6). Consistently, transgenic Arabidopsis plants expressing CPK5 T98A in a *cpk5* mutant background were more resistant to infection with *Pto* DC 3000 compared to plant lines expressing wild type or the T98D variants of CPK5.

**Fig. 6.**
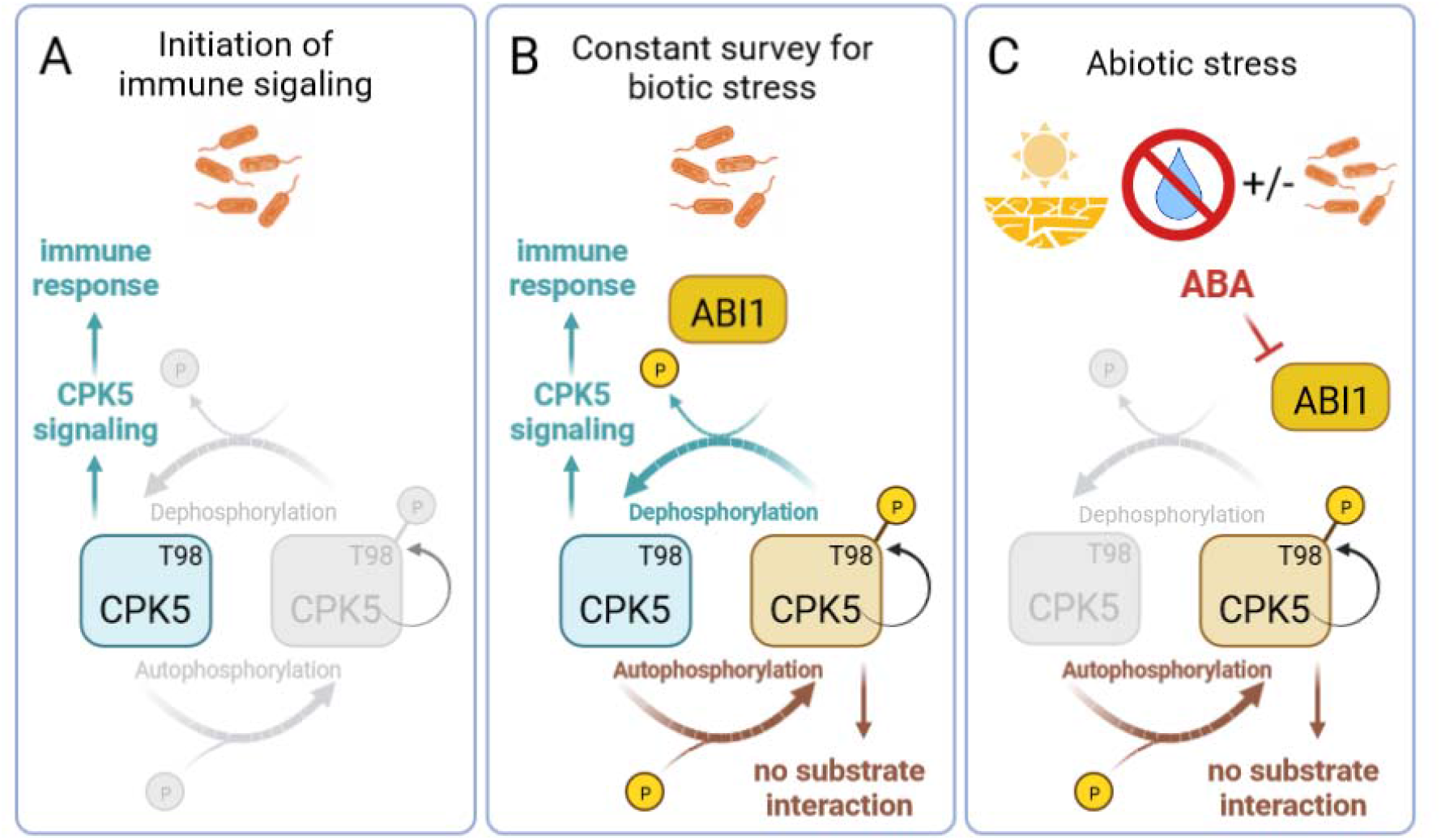
Model for CPK5 T98-mediated phospho-switch between abiotic and biotic stress signaling. (*A*) Pathogen perception by the plant initiates immune signaling, whereby increasing cytoplasmic Ca^2+^ activates CPK5. CPK5 function results in immune responses and defense against the biotic stress. (*B*) Under environmentally non-challenging conditions, auto-phosphorylation of CPK5 at T98 will be balanced through constant de-phosphorylation in the presence of active ABI1 phosphatase. This biochemical switch enables a rapid immune response upon an immediate pathogen threat, and at the same time avoids growth inhibition caused by continuous immune signaling once the pathogen threat is overcome. (*C*) Under abiotic stress conditions ABA signaling is activated and the phosphatase ABI1 is sequestered and inhibited. CPK5 remains phosphorylated at T98, which results in reduced interaction with its substrate(s), and further CPK5-mediated immune signaling is blocked. Even in case of an ongoing pathogen challenge the plant prioritizes in favor of abiotic stress tolerance.

Whereas our research focused on phosphatase ABI1 we cannot rule out contributions of other PP2C phosphatases in the cross-talk between abiotic and biotic immune signaling. In particular not, as we showed that the lack of PP2C phosphatases *HAB1-1* and *PP2CA-1*, both implicated in ABA signaling, compromised CPK5-mediated defense gene expression and resistance to bacterial pathogens (Fig. 2, Fig. S3).

Our focus on phospho-sites in the CPK5 N-terminal and kinase domain has been rationalized by the constitutive active form of CPK5 being sufficient for enhanced ROS production (Fig. 3A). The results on T98 as reversible phospho-switch controlling kinase-substrate interaction justified this choice. However, post-translational modifications by phosphorylation comprise the entire CPK5 protein shown by us and others (57). So, it is already evident from our study that ABI1 is unable to dephosphorylate CPK5 T100 (Fig. 4A). Whether the biochemical context of this distinct site, a phospho-code encompassing multiple phosphorylation sites, hierarchical orders (for example T98 plus S100), or different phosphatases are required remains to be shown. It is thus conceivable that *in planta* several phosphatases (and kinases) with different site-specificities are responsible to regulate the enzyme and its biological pathway (58). This interpretation of multiple joint enzymes is corroborated by our gain-of-function approach employing protein variants ABI1-1 and ABI1-11, which carry the G/D and G/A amino acid substitutions analogous to respective described (semi-)dominant alleles (52–54). Interestingly, enhanced bacterial pathogen resistance has been reported for plants carrying the *abi1-1* semi-dominant allele compared to its respective Ler wild type (44). In our experiments in Col-0 no altered resistance to pathogen *Pto* DC3000 infection occurred in the presence of the *abi1-1* allele (Fig. S5B). In transgenic plants that express active protein phosphatase variant ABI1-11 together with CPK5-YFP no statistic significant additional gain of pathogen resistance to bacterial infection is observed compared to the expression of CPK5-YFP alone (Fig. 3D). This may be explained by the fact that line CPK5-YFP #7 already shows constitutive pathogen resistance, and further enhancement would lead to lethality of plants, as we have observed upon co-expression of the stronger ABI1-1 allele with CPK5-YFP.

Biochemical (auto-)phosphorylation of CDPKs, including CPK5, has often been correlated and interpreted as an activation step, because of its temporal occurrence after stress exposure, for example by flg22 treatment of plants or cells (12–14, 59, 60). In particular, flg22-induced parallel increase in phosphorylation signals *in-gel* kinase assays and a shift to slower migrating enzyme forms due to multiple phospho-groups in western blots was reported for CPK5 (12, 61). Regulation of CDPKs by phosphorylation in addition to Ca^2+^ has been considered as a mechanism to fine-tune kinase activation in terms of strength and duration (18, 20). Phospho-proteomic studies had indicated a rapid de-phosphorylation subsequent to ABA treatment of plants at CPK5 S552 close to the protein’s C-terminus (57).

T98 is located at the N-terminal part of the CPK5 kinase domain. Interestingly, T98A or T98D substitutions confer no difference in biochemical *in-vitro* kinase assays, when using recombinant protein and short peptide substrates (Fig. S8). This does not rule out that *in planta* the phosphorylation status at CPK5 T98 alters the three-dimensional CPK5 protein conformation or protein assembly at a specific sub-cellular region (for example for RBOHD at the plasma membrane), which then facilitates or restricts the interaction of the kinase with its substrates. Such interpretation is consistent, with our data showing under prolonged ABA treatment of plants a reduced CPK5 *in-gel* kinase phosphorylation signal, whereas the CPK5 protein remains present and in its low migrating, post-translationally modified form (Fig. 1H). It is also consistent with our data exemplified with RBOHD as a known CPK5 *in-vivo* substrate, where a strong interaction occurred with CPK5 wild type and T98A variants but weaker interaction with the T98D variant (Fig. 5 D-E, Fig. S9). An alteration of CDPK substrate specificity due to its auto-phosphorylation status has been reported for tobacco *Nt*CDPK1, where phosphorylation at pSer-6 (and pThr-21) within the N-terminal variable domain causes reduced binding affinity to its substrate (62). Employment of a phospho-code at a CDPK has recently been deciphered for *At*CPK28, in which a single phosphorylation site (S318) displays functional diversity in two entirely different signaling programs of immune homeostasis in PTI and stem elongation in plant development (58, 63). In CPK28 the phosphorylation site S318 is located within the kinase domain and its phosphorylation renders the enzyme more sensitive to low Ca^2+^ concentrations, priming a more rapid response upon immune stimulation. Yet, S318 phosphorylation is dispensable for stem development (63).

Our data uncover a link between abiotic ABA signaling and inhibition of biotic immune signaling via reversible phosphorylation of CPK5 as a representative of the calcium-regulated signaling pathway. Interestingly, ABA was reported to induce the expression of PP2C phosphatase genes *HAI1*, *HAI2*, and *HAI3*, and respective phosphatase proteins interact with, de-phosphorylate, and inactivate MAP kinases MPK3 and MPK6, which results in immune suppression via the MAP kinase pathway (49).

In guard cells, ABA signal transduction triggers stomatal closure by employing Ca^2+^-dependent and Ca^2+^-independent pathways, which merge at S-type anion channel (SLAC) phosphorylation and activation. Interestingly, a SA-induced stomatal closure has been reported, which is compromised in guard cells of *cpk3* and *cpk6* mutant plants (37), placing SA upstream of CDPK function and independent of ABA. Further data discuss calcium-sensor kinases including CPK6, CPK21 or CPK23, which contribute to ABA-induced and ion channel-mediated stomata control. These enzymes harbor a significant higher Kd50 [Ca^2+^] of 186 nM (CPK6), and 277 nM (CPK21) than what we observe for CPK5 Kd50 [Ca^2+^] of 100 nM (16, 30). The low CPK5 value may explain why ectopic/heterologous expression of CPK5 in protoplasts or oocytes may lead to similar results as observed for its closest homolog, CPK6, in terms of SLAC1-activation (64). Thus, in guard cells CPK-signaling may contribute to pre-invasive immunity.

In mesophyll cells and in the context of post-invasive long-term immunity, our data link PP2C phosphatases with antagonistic ABA- and CPK5-controlled signaling. ABA exposure triggers the abolition of the negative regulation in ABA signaling, blocking PP2C phosphatase activity as paralleled in respective genetic *pp2c* knockout mutant backgrounds. ABA treatment reverses and inhibits CPK5 signaling-induced immune responses. These include a reset of defense gene (*PR1*, *ICS1*, and *NHL10*) expression (Fig. 1 E-G), reduced immune responses (ROS, Fig. S3D), and ultimately a loss of pathogen resistance against *Pto* DC3000 (Fig. 2C) despite the presence of (biochemically activated, Ca^2+^ bound) CPK5-YFP protein.

## Conclusion

We uncovered reversible phosphorylation at T98 of CPK5 as a rapid biochemical mechanism to disconnect kinase substrate interaction and interrupt CPK5 biological function in immune signaling. This phospho-switch links ABA/ABI1-mediated abiotic signaling and CPK5-mediated SA-dependent immune responses. Our data not only close a mechanistic gap between these antagonistic pathways. The reciprocal control of CPK5 function as a central hub for plant immunity also allows the plant to prioritize in favor of abiotic stress tolerance despite of an ongoing biotic challenge of pathogen pressure (Fig. 6).

## Material and Methods

Detailed descriptions of materials and methods used in this study, including plant growth and treatment conditions, pathogen inoculations, protoplast isolation, protein kinase analyses, gene expression analyses, phytohormone analysis and phosphoproteomics, as well as statistics, are provided in SI Appendix, Materials and Methods. All primers used are listed in SI Appendix, Table S1.

## Data Availability

All study data are included in the article and/or SI Appendix.

## Acknowledgements

We thank Ruth Lintermann for generation of CPK5 amino acid substitution variants, Tobias Lortzing for phytohormone measurements, and Fabian Sylvester for kinase assays with peptide substrates. Plant lines carrying mutations in *PP2C* phosphatase genes (*abi1-1*, *abi1-2*, *hab1-1*, *pp2ca-1*) were kindly provided by Jörg Kudla (University of Münster) and Erwin Grill (Technical University of Munich). This research was supported by Deutsche Forschungsgemeinschaft (DFG) within Collaborative Research Centre SFB973 to TR as well as core institutional funding (Leibniz Foundation).

## Author contributions

HS and TR designed the research. HS, JB, XJ, AL, BC, WH and SM performed the experiments and analyzed the data. HS and TR wrote the manuscript.

---

**Fig. S1.**
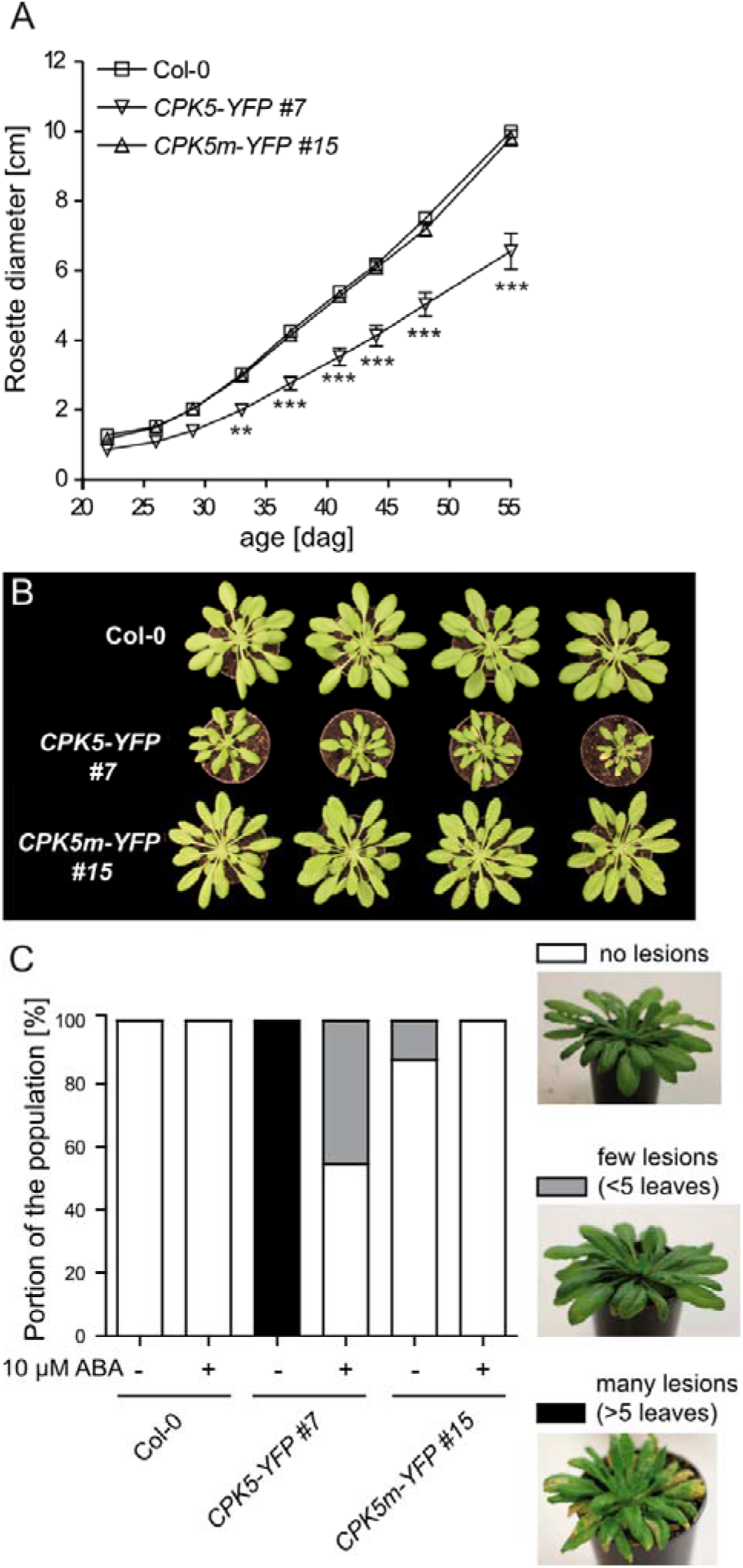
Growth and lesion mimic phenotype of CPK5-YFP overexpression lines. (*A*) Rosette diameter (n = 10-14) and (*B*) phenotype of 8 week old plants of the lines Col-0, *CPK5-YFP #7* and *CPK5m-YFP #15*. Values represent mean ± SE of n biological replicates. Data in (*A*) were evaluated for statistical differences using two-way ANOVA (Bonferroni post-test, significant differences ** = p < 0.01 and *** = p < 0.001). (*C*) Evaluation of lesions of 70 days old plants of the lines Col-0, *CPK5-YFP #7* and *CPK5m-YFP #15* as in (*A*) + (*B*) with examples of categories for evaluation of lesions.

**Fig. S2.**
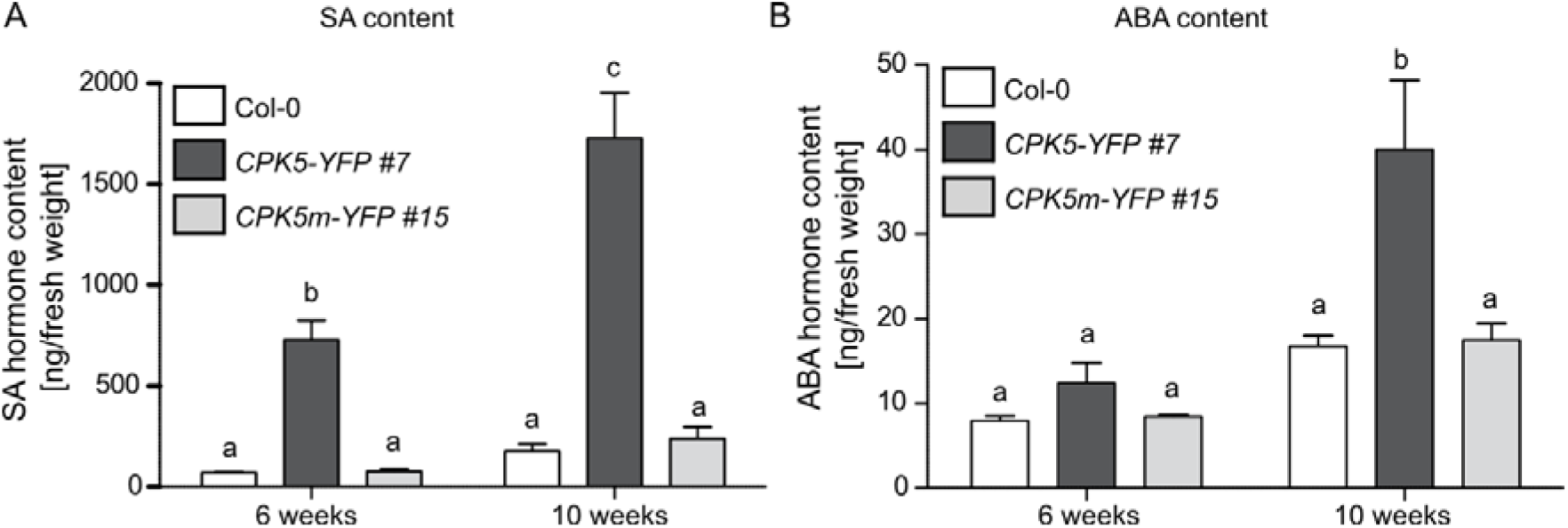
Basal phytohormone contents in *CPK5-YFP* lines. Basal SA-(*A*) and ABA-levels (*B*) in untreated 6 and 10 weeks-old plants of the lines Col-0, *CPK5-YFP #7* und *CPK5m-YFP #15* (n = 5-7) measured via UPLC-TOF-MS/MS. Values represent mean ± SE of n biological replicates. Data were evaluated for statistical differences using two-way ANOVA (Bonferroni post-test against all groups, p < 0.05, significant differences are indicated with different letters).

**Fig. S3.**
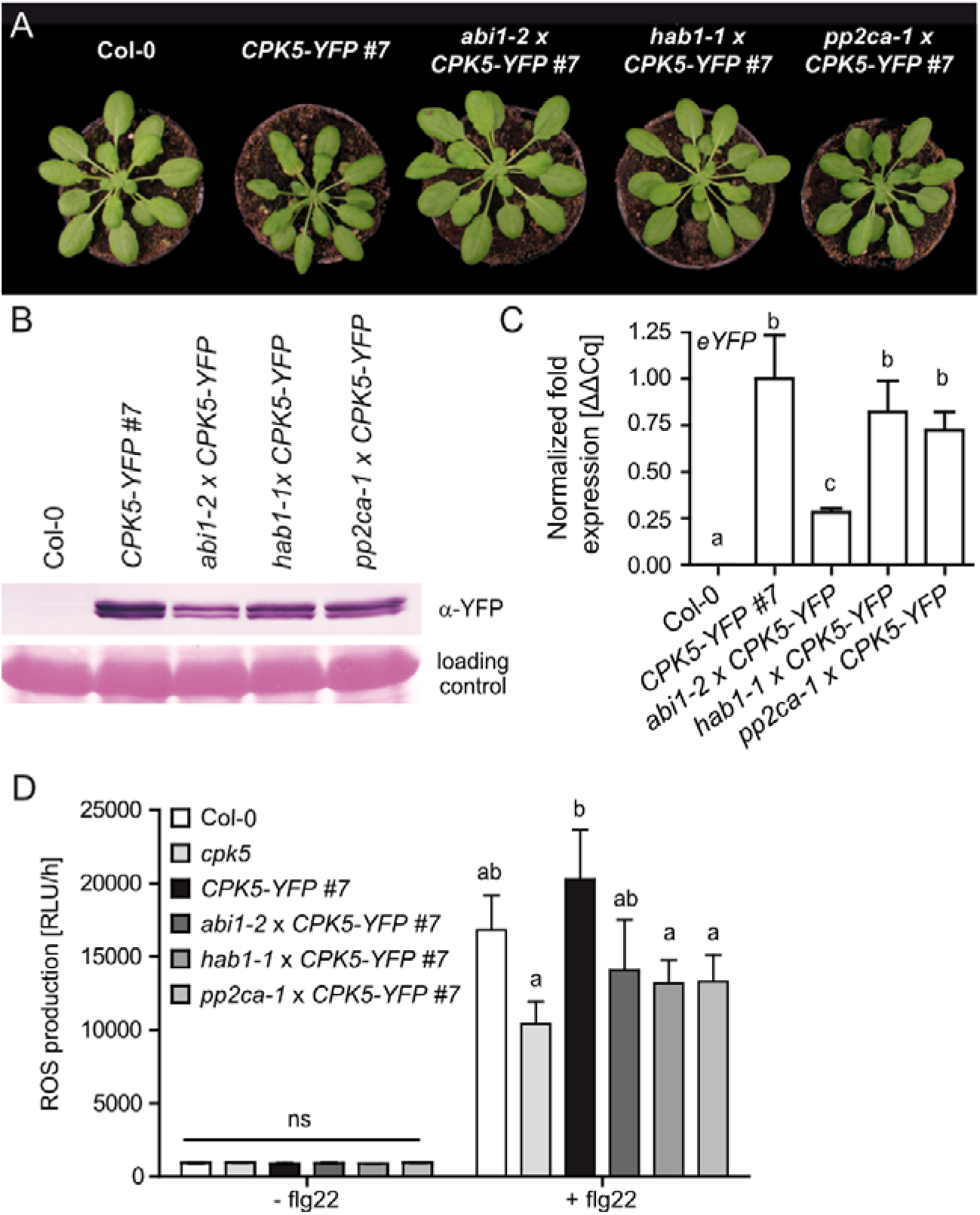
CPK5-YFP phenotype is reverted in different *pp2c* knock out backgrounds. (*A*) Phenotype of 6 week old plants of the lines Col-0, *CPK5-YFP #7* and the crossing lines *abi1-2 x CPK5-YFP #7, hab1-1 x CPK5-YFP #7* and *pp2ca-1 x CPK5-YFP #7*. (*B*) Western blot analysis of CPK5-YFP expression in 3 week old plants as in (*A*) via detection of YFP. (*C*) Transcript levels of YFP in 6 week old plants as mentioned in (*A*) (mean ± SE, n = 5-11). YFP expression in *CPK5-YFP # 7* was set to 1. (*D*) ROS levels measured with a luminol-based assay in absence or presence of 200 nM flg22 over 1h in the lines Col-0, *cpk5, CPK5-YFP #7* and crossing lines as in (*A*). Values represent additive ROS production over 1 hour and depict mean ± SD of 4-12 biological replicates. Data were evaluated for statistical differences using one-way (*C*) or two-way ANOVA (*D*) with Bonferroni post-tests comparing all groups, p < 0.05, significantly different groups were assigned different letters.

**Fig. S4.**
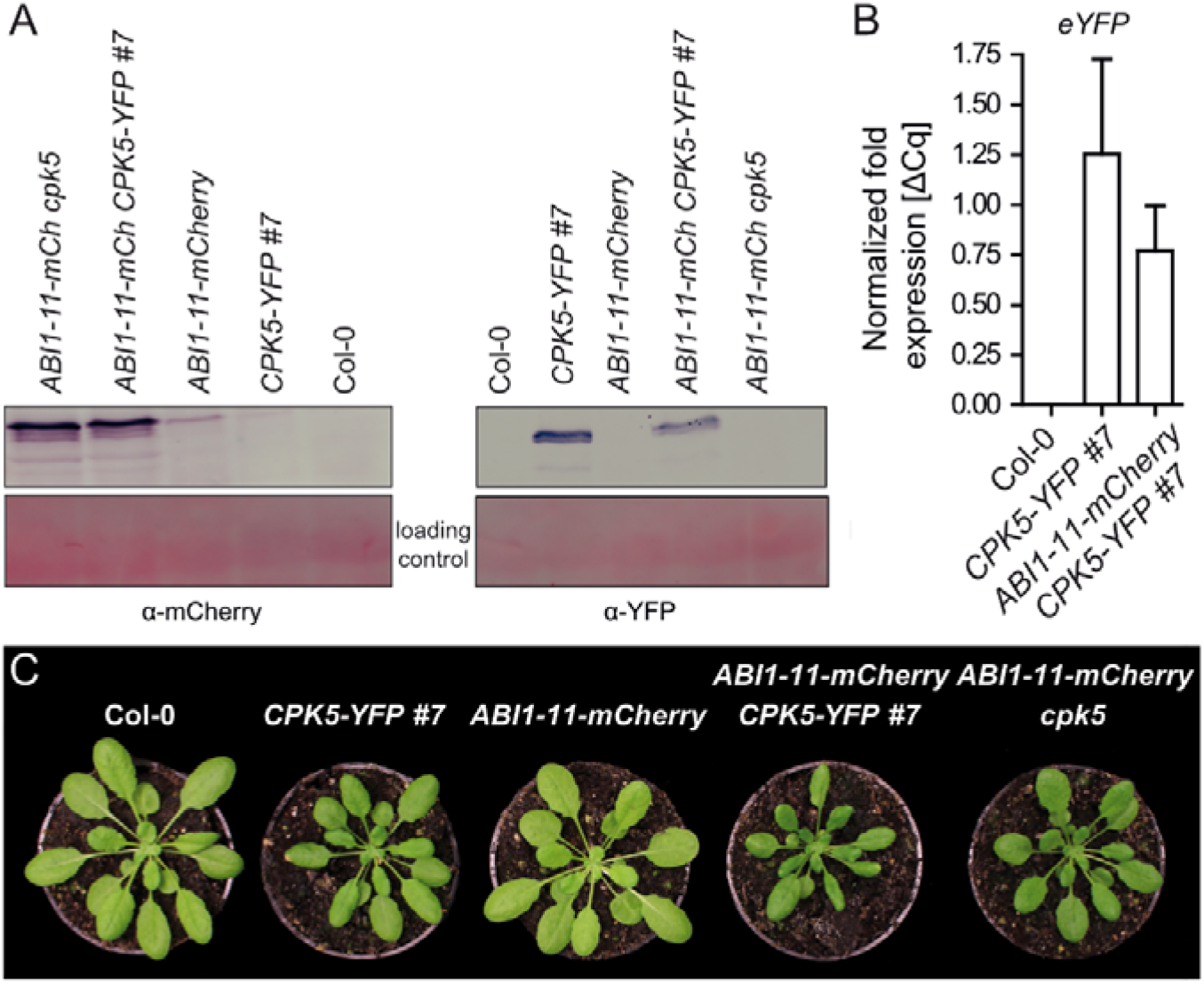
Phenotype of ABI1-11-mCherry lines. (*A*) Western blot analysis of protein expression in the lines Col-0, *CPK5-YFP #7, ABI1-11-mCherry, ABI1-11-mCherry CPK5-YFP #7* and *ABI1-11-mCherry cpk5* via detection of mCherry and YFP. (*B*) Transcript levels of YFP in Col-0, *CPK5-YFP #7* and *ABI1-11-mCherry CPK5-YFP #7* in 6 weeks-old plants (bars represent mean ± SE n = 4). (*C*) Phenotype of 6 week old plants of the lines as mentioned in (*A*).

**Fig. S5.**
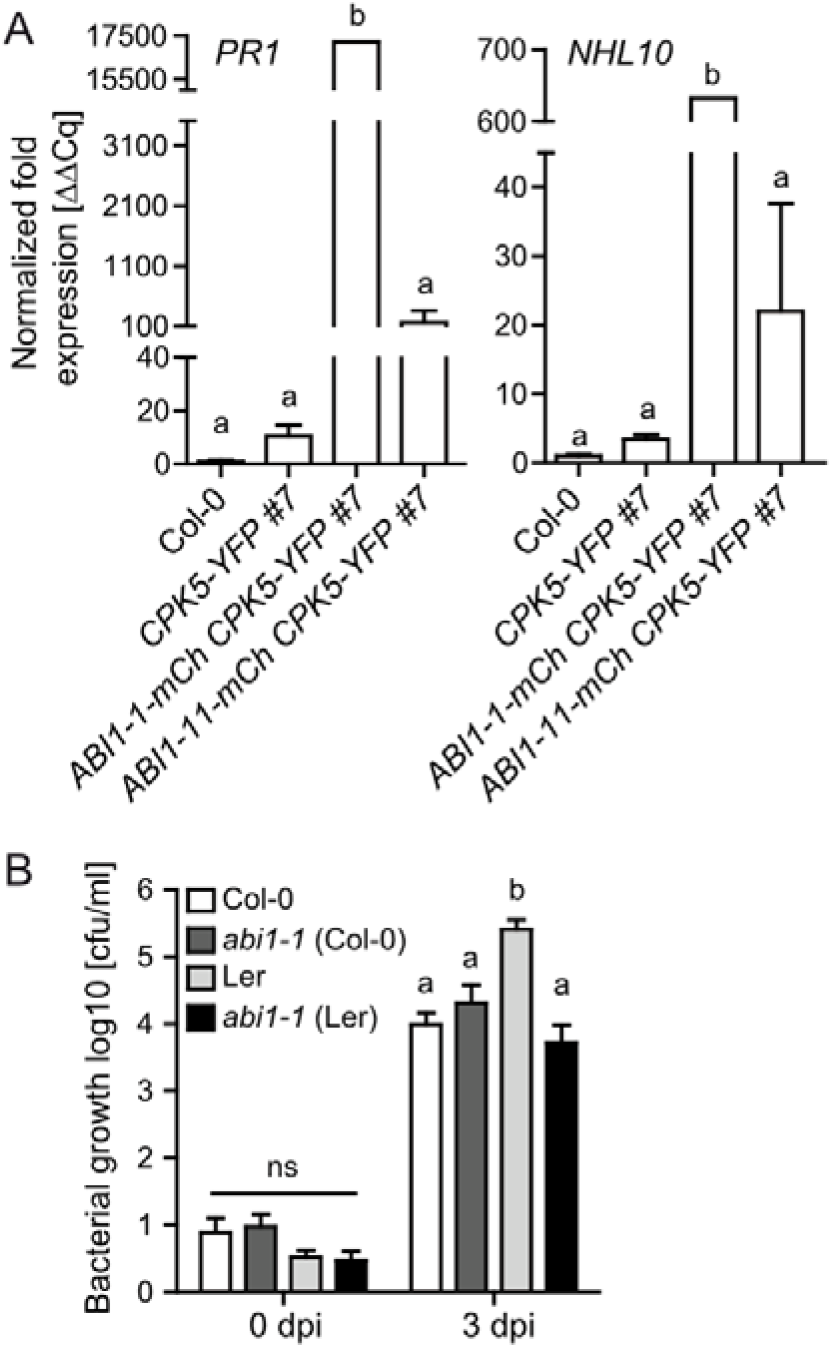
Characterization of ABI1-1-mCherry and *abi1-1* transgenic plants. (*A*) Basal transcript levels of *PR1* and *NHL10* in 6 week old plants of the lines Col-0, *CPK5-YFP #7, ABI1-1-mCherry CPK5-YFP #7* and *ABI1-11-mCherry CPK5-YFP #7* (n = 1-10). (*B*) Bacterial growth of *Pto* DC3000 tested directly and 3 days after infection in knock-out mutants of *ABI1* in different wild type backgrounds (n = 9-12). Values represent mean ± SD of n biological replicates. Data were evaluated for statistical differences using one-way (*A*) or two-way ANOVA (*B*) with Bonferroni post-tests comparing all groups, p < 0.05, significantly different groups were assigned different letters.

**Fig. S6.**
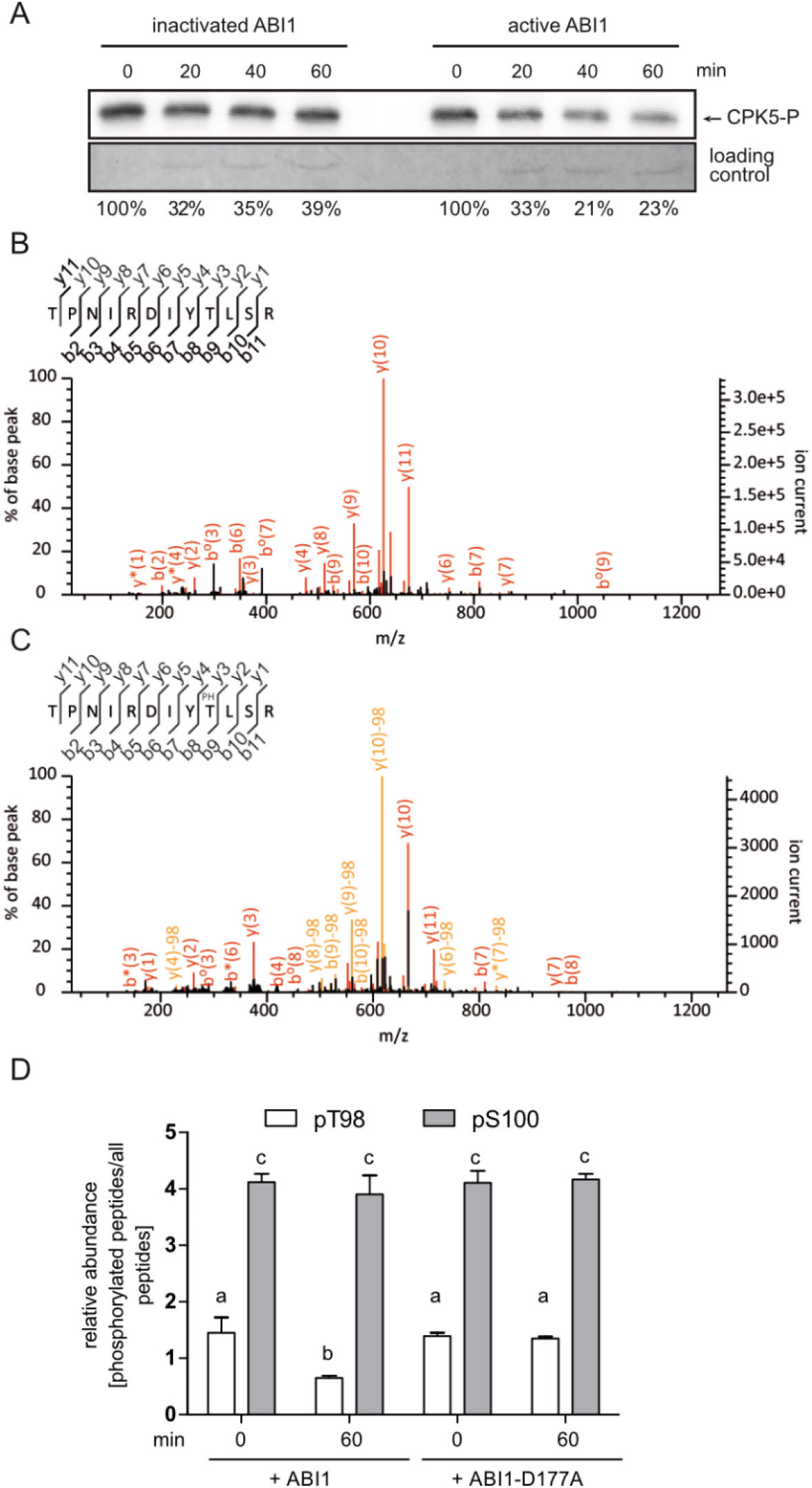
ABI-dependent phosphorylation of CPK5. (*A) In-vitro* kinase assay with 200 ng recombinant CPK5-StrepII and subsequent dephosphorylation with 100 ng recombinant ABI1-StrepII (active or heat-inactivated). CPK5 was inactivated with K252-a previous to coincubation with ABI1. Percentage corresponds to amount of auto-phosphorylated kinase after incubation with ABI1 phosphatase normalized to the loading control. (*B, C*) MS spectra corresponding to peptides of T98 (*B*) and pT98 (*C*) identified in Fig. 4A. (*D*) Targeted MS-analysis of the T98 and S100 phospho-sites. Recombinant, auto-phosphorylated CPK5-StrepII was incubated with active ABI-Strep or catalytically inactive ABI-D177A-Strep for 0 or 60 min. For quantification of phosphorylation, the summed area under the curve (AUC) of fragment ions was used and relative abundance of phosphorylated peptides compared to all peptides spanning the modification site were calculated. pT98 data from this experiment are also shown in Fig. 3B. Values represent mean ± SD of 2 biological replicates. Data were evaluated for statistical differences using two-way ANOVA (Bonferroni’s post-test, p < 0.05). Significantly different groups were assigned different letters.

**Fig. S7.**
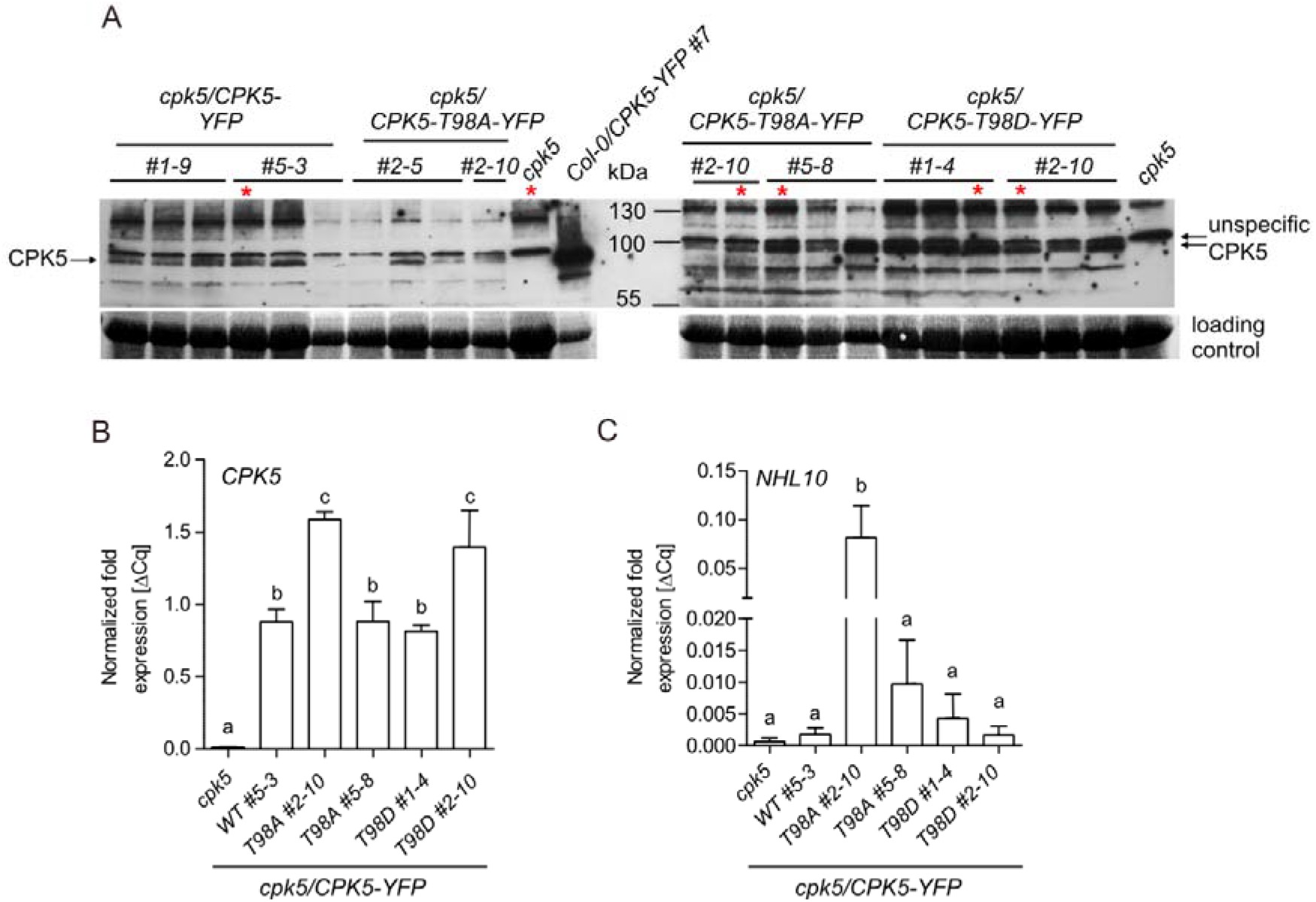
Characterization of transgenic Arabidopsis lines carrying CPK5-T98 phospho-variants in a *cpk5* mutant background. (*A*) Western Blot of protein expression levels of wild-type, phosphomutant (T98A) or phosphomimic (T98D) variant lines of CPK5 in a *cpk5* mutant background for comparison to *cpk5* mutant (no band) and Col-0/CPK5-YFP #7 line (much higher expression despite loading of 1/5^th^ of crude extract). Red stars depict lines used for further experiments and constitute the same Western blot bands used in Fig. 5B. (*B-C*) Basal transcript levels of *CPK5 (B*) and marker gene *NHL10 (C*) in 6 week old plants of T98 phospho variant lines (n=5). Data were evaluated for statistical differences using one-way ANOVA (Bonferroni’s post-test, p < 0.05). Significantly different groups were assigned different letters.

**Fig. S8.**
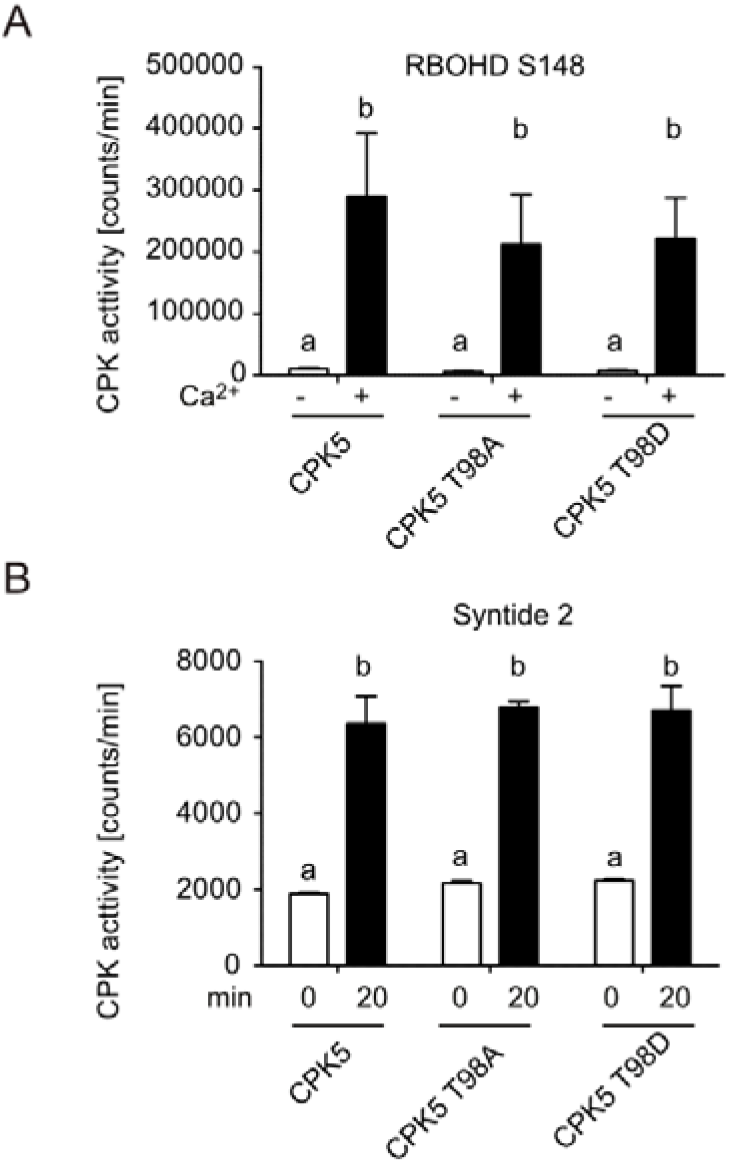
Biochemical kinase activity toward peptide substrates is unaltered in CPK5-T98 phospho-variants. Affinity-purified recombinant protein CPK5-YFP-StrepII (*A*) or CPK5-StrepII (*B*) was analyzed in an *in-vitro* kinase assay towards 10 μM substrate peptide of respiratory burst oxidase homolog protein D (RBOHD) encompassing S148 (10-amino-acid peptide) (*A*) or Syntide 2 (*B*). Phosphorylation activity in the absence or presence of Ca^2+^ was monitored within 20 min. Values are means ± SEM of two technical replicates. Data were evaluated for statistical differences using two-way ANOVA (Bonferroni’s post-test, p < 0.05). Significantly different groups were assigned different letters.

**Fig. S9.**
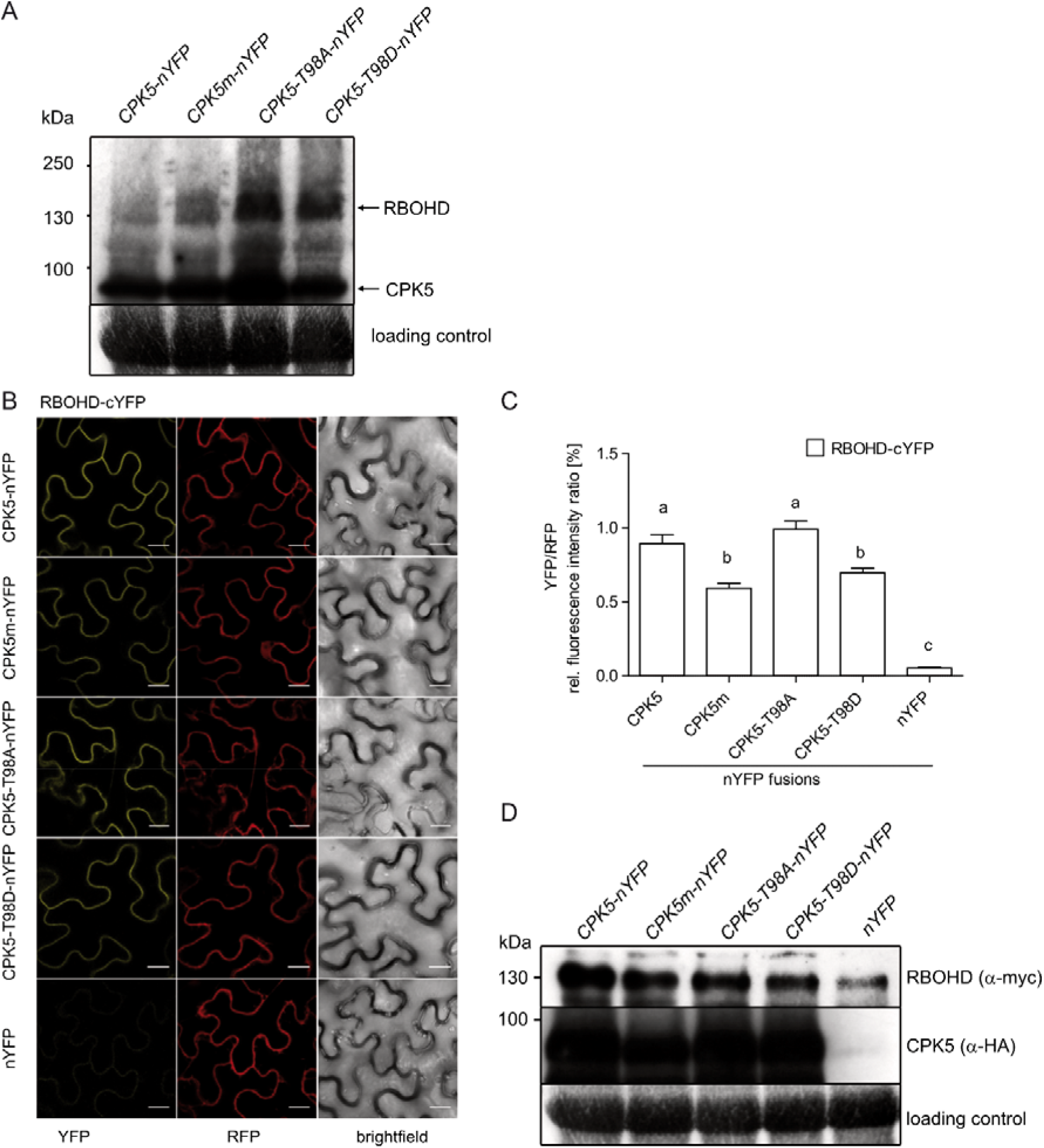
Evaluation of *in-vivo* CPK5 and RBOHD interaction by BiFC in *N. benthamiana*. (*A*) Confirmation of CPK5 and RBOHD protein expression from BiFC experiment shown in Fig. 5D by Western Blot. (*B-D*) Ratiometric BiFC experiment of *in-vivo* CPK5 and RBOHD interaction in *N. benthamiana* using a one-vector system with RFP as internal expression control. (*C*) Quantification of fluorescence signals from YFP and RFP channels via ImageJ with RFP fluorescence signal as a normalization factor. Values represent mean ± SD of n biological replicates (n = 20). Quantification data were evaluated for statistical differences using one-way ANOVA (Bonferroni’s post-test, p < 0.05). Significantly different groups were assigned different letters. Scale bar = 20 μM. (*D*) CPK5 and RBOHD protein expression of BiFC experiment in (*B,C*) confirmed by Western Blot.

## Supplementary Information Appendix

### SI Appendix Materials and Methods

#### Plant material

*Arabidopsis thaliana* plants were grown on soil with 8 h light/16 h dark cycle, 20 °C and 60 % relative humidity. T-DNA insertion lines *abi1-2* (SALK_072009), *hab1-1* (SALK_02104), *pp2ca-1* (SALK_28132), *abi2-2* (SALK_15166) as well as *abi1-1* (Col-0 background, dominant mutant) were kindly provided by J. Kudla (WWU Münster). *abi1-1* (Ler background, dominant mutant) and the Ler ecotype were kindly provided by E. Grill (TU München). *cpk5* (SAIL_657_C06) was obtained from Nottingham Arabidopsis Stock Center (NASC). Transgenic overexpression lines *CPK5-YFP #7* and *CPK5m-YFP #15* (kinase-deficient variant carrying single amino acid substitution CPK5-D221A) were generated previously (12). *N. benthamiana* wildtype plants were grown on soil in a greenhouse with 16 h light/8 h dark cycle.

#### Generation of constructs and transgenic lines

Generation of the ABI1 point mutation variants ABI1-1 (G180D) and ABI1-11 (G181S) and inactive ABID177A (65) and CPK5 phospho-site mutants, T98A/D, were performed using PCR mutagenesis (for primers see Table S1). ABI1-1 and ABI1-11 were put into the binary vector pXCSnpt-p35S::mCherry. CPK5 and phospho-site mutants T98A/D were cloned into pDONR221 (Invitrogen) and put into pXCSG-p35S::YFP by LR-Gateway recombination. Overexpression lines were generated using the floral dipping method (66) in the respective background lines (Col-0, *cpk5, CPK5-YFP #7*). For 2-vector BiFC experiments, RBOHD and CPK5 or CPK5 phospho-site variants T98A and T98D were cloned into binary vectors pUBC-cYFP or pUBC-nYFP by Gateway cloning, respectively. For the ratiometric BiFC method, RBOHD and CPK5 or CPK5 phospho-site variants were cloned into pDONR221-P1P4 and pDONR221-P3P2 (Invitrogen), respectively. Both pENTRY clones were put into a binary vector pBiFCt-2in1-CC (56).

#### *In-gel* kinase and *in-vitro* kinase and phosphatase assays

*In-gel* kinase assays for assessing *in-vivo* kinase activity in crude extracts and immunoblot analysis of proteins were performed as described in detail previously (12, 67). CDPK *in-gel* kinase assay: Protoplasts were harvested by centrifugation and frozen in liquid nitrogen. Protoplast pellets or frozen Arabidopsis leaf material samples were resuspended by vortexing in extraction buffer (100 mM Tris-HCl (pH 8.0), 200 mM NaCl, 20 mM DTT, 10 mM NaF, 10 mM NaVO4, 10 mM β-glycerole-phosphate, 0.5 mM AEBSF, 100 μg/mL avidin and 1 × protease inhibitor mixture (Sigma)). Crude extracts were centrifuged and proteins were separated by SDS-PAGE containing 0.25 mg/ml MBP or Histone III-S (Sigma). The gel was washed two times 1 h in wash-buffer (25 mM Tris-HCl, pH 7.5; 0.5 mM DTT; 5 mM NaF; 0.1 mM Na_3_VO_4_; 0.5 mg/ml BSA; 0.1 % Triton X-100). Protein renaturation was performed by incubating the gel with renaturation buffer (25 mM Tris-HCl, pH 7.5; 0.5 mM DTT; 5 mM NaF; 0.1 mM Na_3_VO_4_) for 1 h at room temperature, overnight at 4 °C and 1 h at room temperature. After equilibration of the gel for 30 min in the reaction buffer (25 mM Tris-HCl, pH 7.5; 1 mM DTT; 0.1 mM Na_3_VO_4_; 12 mM MgCl_2_; 1 mM CaCl_2_), the kinase reaction was performed for 1.5 h in the reaction buffer with 0.1 nM ATP and 50 μCi [γ-32P]-ATP. The reaction was stopped and washed 6 times with 5 % TCA and 1 % phosphoric acid for 4 h. The gel was dried and visualized by autoradiography.

*In-vitro* kinase assays were performed with affinity purified *At*CPK5 protein (StrepII or YFP-StrepII) expressed in *N. benthamiana*. Strep-Tactin Macroprep-bound kinase was resuspended with Buffer E (50 mM HEPES, pH 7.4; 2 mM DTT; 0.1 mM EDTA). 5 μl slurry was mixed with 20 μl Buffer E and 5 μl radiolabeled reaction mix (60 mM MgCl2; 60 μM CaCl_2_; 60 μM ATP; 60 μM Syntide 2 or 60 μM *At*RBOHD peptide (aa 143-152); 3 μCi [γ-32P]-ATP). As negative control, CaCl_2_ was replaced by 12 mM EGTA. The reaction was performed for 20 min at RT and stopped with 3 μl 10% phosphoric acid. 20 μl of the reaction was spotted on P81 anion-exchange paper, washed 4 times with 1 % phosphoric acid and radioactivity was determined using scintillation mixture.

*In-vitro* phosphatase assays were carried out with recombinant CPK5-Strep, ABI1-Strep and ABI1-D177A-Strep purified from *E. coli*. In radiolabelled dephosphorylation assays, an initial step of CPK5 kinase autophosphorylation for 15 minutes was carried out in buffer E as described above, and the reaction was stopped with kinase inhibitor K252 for 15 min. ABI1 or heat-inactivated ABI1 phosphatase as negative control was added to the reaction mix and dephosphorylation was stopped by heating for 5 min at 95 C in 5x SDS buffer. Samples were analyzed immediately or stored at-20 °C before SDS-PAGE and autoradiography. In unlabelled assays for proteomic analysis, the initial step of CPK5 kinase autophosphorylation was carried out with 10 μl reaction mix (10 mM MgCl, 10 μM ATP, 5 μM CaCl2) and 40 μl buffer (30 mM Tris pH 8,0, 180 mM KCl). The reaction was stopped by addition of 40 μl stop buffer (30 mM Tris pH8,0, 180 mM KC, 10 mM MgCl, 3,4 μM kinase inhibitors K252, 10 mM Biotin). After addition of ABI1 or ABI1-D177A for the specified amount of time, dephosphorylation was terminated by heating for 5 min at 95 C in 5 x SDS buffer. Samples were analyzed immediately or stored at-20 °C before SDS-PAGE and subsequent MS analysis.

#### Measurement of transcript levels by qRT-PCR analysis

RNA was extracted from ground leaf tissue using the Trizol method. 2 μg of total RNA was treated with RNase-free DNase (Fermentas) and mRNA was reverse transcribed with SuperscriptIII SuperMix (Invitrogen) according to the manufacturer’s protocols. Real-time quantitative PCR analysis was performed according to the instructions of Power SYBR Green PCR Master Mix (Applied Biosystems) using the CFX96 system (Bio-Rad). Post-amplification dissociation curves were used for evaluation of amplification specificity. Gene expression was quantified using *ACTIN2* (At3G18780) or *PP2A* (At1G13320) and primer sequences are listed in Table S1.

#### ROS measurement

Production of reactive oxygen species (ROS) was monitored using a leaf disc assay described previously (12). In short, reactive oxygen species production was monitored using a luminol-dependent assay. 0.5 cm diameter leaf discs were floated on 100 μl H_2_O in a 96-well plate overnight. On the following day, 100 μl assay solution (final concentration: 0.034 mg/ml luminol and 0.02 mg/ml horse radish peroxidase) with or without 200 nM flg22 was added to the leaf discs and luminescence was immediately measured over one hour. Values represent accumulation of total ROS measured over one hour. ROS production in *A. thaliana* was tested as flg22-induced oxidative burst. In *N. benthamiana*, ROS was measured in the absence of flg22 3 days after *A. tumefaciens-*mediated transformation with constitutively active CPK5-VK constructs.

#### ABA treatment

Plants were treated with ABA solutions by spraying whole rosettes with a manual aerosol until all leaves were moistened. Intermediate-term (14 days) effects of ABA on CPK5-YFP activity and lesion phenotype were tested after spray treatment of plants with 3 μM ABA (in 30 μM MES, pH 5.7) every second day. Long-term effects of ABA treatment on the CPK5-YFP lesion phenotype were monitored over a growth period of 10 weeks under continuous treatment with 10 μM ABA (in 30 μM MES, pH 5.7) every second day.

#### Bacterial growth

Growth of *P. syringae* pv. *tomato* DC3000 (*Pto* DC3000) was monitored as described before (12). Summarized, 6-weeks old plants were syringe infiltrated with 10^4^ cfu/ml *Pto* DC3000 in 10 mM MgCl_2_. Bacterial growth was tested three days after inoculation by serial dilution plating of leaf disc extracts.

#### Phytohormone analysis with UPLC-TOF-MS/MS

Plant extracts were prepared after a modified method described previously (68). 1 ml extraction buffer (100% ethyl acetate, deuterized internal standard mix), 500 mg Matrix D (*Fast-Prep Homogenizer*) and 2 steal beads were added to 200-300 mg plant material and mixed at RT for 1 min at 5 m/sec in 24/2 FastPrep^®^-24 (MP Biomedicals) followed by 30 sec incubation on ice. 1 ml supernatant (after centrifugation at 16.000 x g, 4 °C, 10 min) was vacuum-concentrated. The remaining pellet was dissolved in 1 ml extraction buffer, centrifuged twice and the supernatant was combined with the first one and again concentrated for 20-30 min. The pellet was dissolved in 400 μl Re-eluation buffer (70 % Methanol, 0.1 formaldehyde) and mixed for 10 min at RT. Samples were loaded on a C18 column (Acquinity UPLC BEH-C18, length 5 cm, ID 2.1 mm, particle size 1,7 μm) for hormone separation at 4 °C. Elution was started with constant 30 % eluent B (Methanol with 0.1 formic acid, eluent A was water with 0.1 % formic acid) followed by a linear gradient to 90 % of eluent B for 3.5 min and constant 90 % eluent B for another 3.5 min. Eluent B was decreased to 30 % for 1 min and kept at 30 % for additional 3 min. Flow rate was 0.25 ml/min with an injection volume of 7 μl per sample. Synapt^®^ G2-S HDMS ESIMS/ MS (Waters) was used for detection of phytohormones in negative ion mode. Electrospray ionization conditions were capillary tension 2.50 kV, vaporizer 6.0 bar, desolvatation of gas flow rate 500 l/h, source temp 80 °C, desolvatation temp 150 °C. The specific fragment sizes were SA – m/z 93, ABA – m/z 153.

#### Phosphoproteomics

Amino acid residue-specific phosphorylation of AtCPK5 was mapped by liquid chromatography on-line with high resolution accurate mass MS (HR/AM LC-MS). Proteins separated by SDS-PAGE were subjected to *in-gel* tryptic digestion. The resulting peptides were separated using C18 reverse phase chemistry employing a pre-column (EASY column SC001, length 2 cm, ID 100 μm, particle size 5 μm) in line with an EASY column SC200 with a length of 10 cm, an inner diameter (ID) of 75 μm and a particle size of 3 μmon an EASY-nLC II (all from Thermo Fisher Scientific). Peptides were eluted into a Nanospray Flex ion source (Thermo Fisher Scientific) with a 90 min gradient increasing from 5 % to 40 % acetonitrile in ddH_2_O with a flow rate of 300 nl/min and electrosprayed into an OrbitrapVelos Pro mass spectrometer (Thermo Fisher Scientific). The source voltage was set to 1.9 kV, the S Lens RF level to 50 %. The delta multipole offset was −7.00. The AGC target value was set to 1e06 and the maximum injection time (max IT) to 500 ms in the Orbitrap. The parameters were set to 1e04 and 100 ms in the LTQ with an isolation width of 2 Da for precursor isolation and MS/MS scanning. Peptides were analyzed by a targeted data acquisition (TDA) scan strategy with inclusion list to specifically select and isolate *At*CPK5 phosphorylated peptides for MS/MS peptide sequencing. Multi stage activation (MSA) was applied to further fragment ion peaks resulting from neutral loss of the phosphate moiety by dissociation of the high energy phosphate bond to generate b- and y-fragment ion series rich in peptide sequence information.

MS/MS spectra were used to search the TAIR10 database (ftp://ftp.arabidopsis.org, 35394 sequences, 14486974 residues) with the Mascot software v.2.5 linked to Proteome Discoverer v.1.4. The enzyme specificity was set to trypsin and two missed cleavages were tolerated. Carbamidomethylation of cysteine was set as a fixed modification and oxidation of methionine and phosphorylation of serine, threonine and tyrosine as variable modifications. The precursor tolerance was set to 7 ppm and the product ion mass tolerance was set to 0.8 Da. A decoy database search was performed to determine the peptide spectral match (PSM) and peptide identification false discovery rates (FDR). Peptides identifications with an ion score of 24 surpassing the Mascot significance threshold (p<0.05) which corresponded to a peptide identification FDR of < 4.55 % and a PSM FDR of 1.14%, were accepted. A transferred FDR specifically for phosphopeptide PSMs was calculated according to Fu et al. (69) and was 4.47 %. The phosphoRS module was used to specifically map phosphorylation to amino acid residues within the primary structure of phosphopeptides. Measurement from two experiments each of *At*CPK5 with and without ABI1 were concatenated. In each of the concatenated results, the ratio of the number of T98 phosphopeptide spectral matches (#PSM) to the #PSM of non-phosphorylated counterpart peptides was used to quantify site-specific phosphorylation at T98. It was normalized to the fraction of total *At*CPK5 #PSM to total recorded MS2 spectra in each of the concatenated results. This normalized phosphorylation value was further scaled by setting it to 1 for *At*CPK5 phosphorylation without ABI1.

For targeted MS-analysis (parallel reaction monitoring, PRM) distinguishing the T98 and S100 phospho-sites in the same peptide, dried peptides were dissolved in 5% acetonitrile, 0.1% trifluoric acid and injected into an EASY-nLC 1200 liquid chromatography system (Thermo Fisher Scientific). Peptides were separated using liquid chromatography C18 reverse phase chemistry employing a 120 min gradient increasing from 1% to 40% acetonitrile in 0.1% FA, and a flow rate of 250 nL/min. Eluted peptides were electrosprayed on-line into a Fusion Lumos Tribrid mass spectrometer (Thermo Fisher Scientific). The spray voltage was 2.0 kV and the capillary temperature 305°C. A full MS survey scan was carried out with chromatographic peak width set to 15 s, resolution 60,000, automatic gain control (AGC) set to standard and a max injection time (IT) of 100 ms. MS/MS peptide sequencing was performed using a PRM scan strategy (without retention time scheduling) with HCD fragmentation containing 11 target peptide m/z on a list (Supplemental table S2). Top 15 MS/MS scans were acquired in the Orbitrap with resolution 15,000, mass to charge ratios (m/z) between 300 and 2000, AGC target set to 300%, Maximum IT 22 ms, isolation width 1.6 m/z, and normalized collision energy 27%.

Peptides and proteins were identified using the Mascot software v2.7.0 (Matrix Science) linked to Proteome Discoverer v2.1 (Thermo Fisher Scientific) with settings described earlier, except for the following: a precursor ion mass error of 5 ppm and a fragment ion mass error of 0.02 Da were tolerated in searches. For PRM quantification analyses with Skyline (version 20.2.0.343), a spectral library was generated using Mascot search results, applying a cut-off score of 0.95. Ambiguous peptide matches were included, and the library was filtered for peptides spanning the T98 and S100 phosphosites. *.raw files were imported into Skyline and automated fragment ion selection by Skyline was utilized (6 ions/peptide): MS/MS ion trace filtering (centroid mode) and charge states of 1+/2+/3+ for b- and y-ions as well as 2+/3+ for precursor ions. Integration boundaries of peptides were inspected manually and corrected, if necessary. Peptides with truncated peaks or no MS/MS signal were excluded from further analysis. Reports were exported and further processed in MS Excel. For peptide quantification, the summed area under the curve (AUC) of fragment ions was used and relative abundance of phosphorylated peptides compared to all peptides spanning the modification site were calculated.

#### BiFC analysis

Two-vector method: Constructs pUBC-RBOHD-cYFP, pUBC-CPK5-nYFP, pUBC-CPK5m-nYFP, pUBC-CPK5-T98A-nYFP, pUBC-CPK5-T98D-nYFP and pUBC-FLS2-RFP for normalization were transformed into *Agrobacterium tumefaciens* strain GV3101. FLS2-RFP and RBOHD-cYFP was co-infiltrated with different CPK5-nYFP variants into *N. benthamiana*. Images were taken 3 days after infiltration using an LSM780 microscope (Zeiss). Fluorescence signals from YFP and RFP channels were quantified by ImageJ software, and RFP fluorescence signal was used as a normalization factor.

Ratiometric BiFC Method: The binary vector pBiFCt-2in1-CC (56) was used to control for equal expression of cYFP or nYFP fusion proteins as well as RFP as normalization factor within one single vector. Constructs with combinations of RBOHD/CPK5 variants in pBiFCt-2in1-CC ((RBOHD/CPK5), (RBOHD/CPK5mut), (RBOHD/CPK5-T98A), (RBOHD/CPK5-T98D) and (RBOHD/--)) were transformed into *Agrobacterium tumefaciens* strain GV3101. Images were taken 2 days after infiltration using an LSM880 (Zeiss). The fluorescence signals from YFP and RFP channels were quantified by ImageJ software, and RFP fluorescence signal was used as a normalization factor.

**Table S1.**
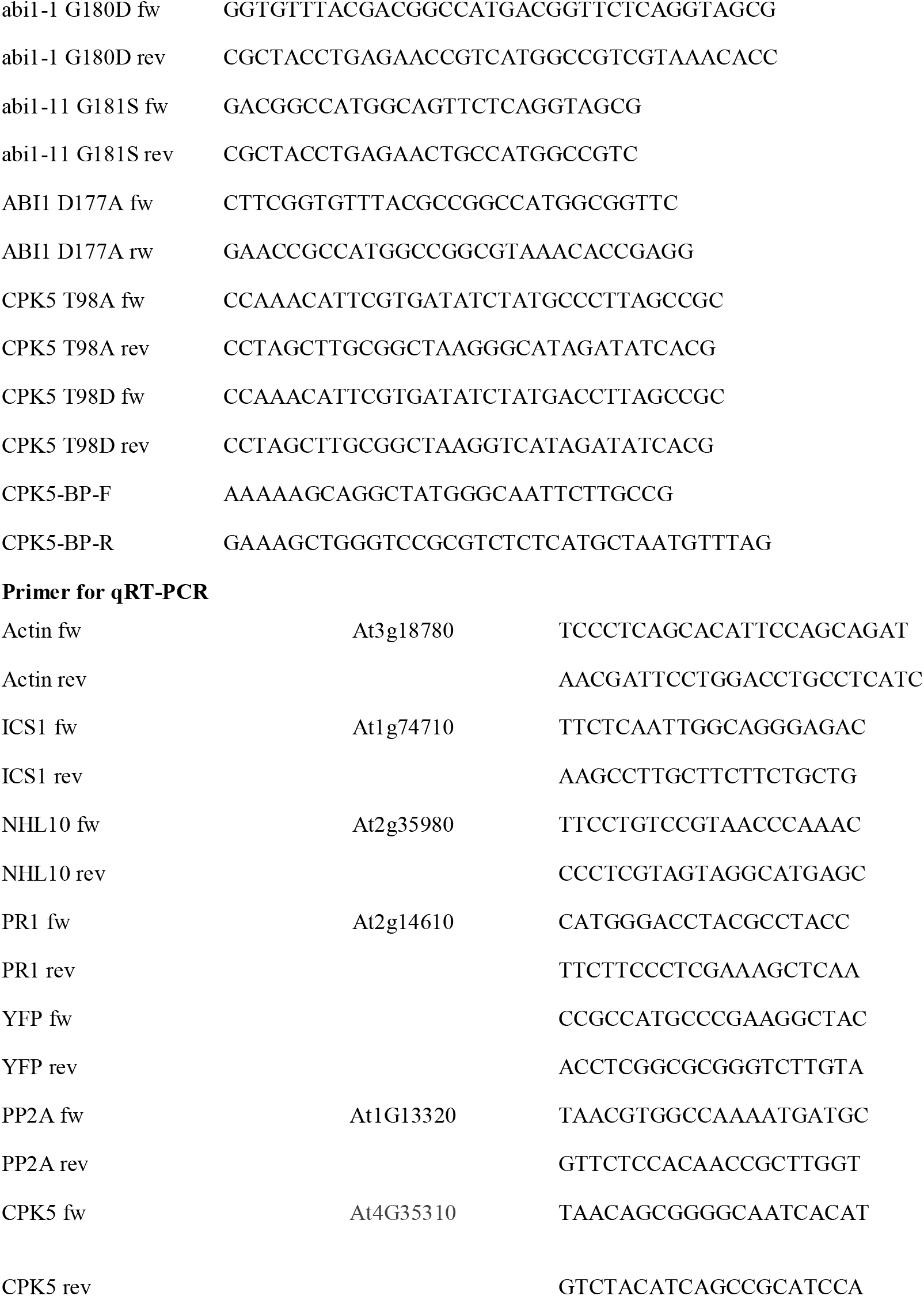
Primer for PCR-Mutagenesis, cloning and qRT-PCR.

**Table S1.** PRM target list (Fusion Lumos)

| Compound | Formula | Adduct | m/z | z |
| --- | --- | --- | --- | --- |
|  |  | (no adduct) | 316.4793 | 3 |
|  |  | (no adduct) | 343.1348 | 3 |
|  |  | (no adduct) | 332.5222 | 3 |
|  |  | (no adduct) | 359.1776 | 3 |
|  |  | (no adduct) | 385.8331 | 3 |
|  |  | (no adduct) | 434.2322 | 2 |
|  |  | (no adduct) | 474.2153 | 2 |
|  |  | (no adduct) | 514.1985 | 2 |
|  |  | (no adduct) | 498.2796 | 2 |
|  |  | (no adduct) | 538.2628 | 2 |
|  |  | (no adduct) | 578.246 | 2 |

## Notes

### Competing Interest Statement

The authors have declared no competing interest.

### Summary of Updates

In the modified version, we performed new experiments to substantiate our argumentation toward the CPK5/ABI1 kinase/phosphatase switch via reversible phosphorylation status at CPK5 T98 site. In addition, we offer new results and a mechanistic explanation on how the T98 auto-phosphorylation site functions by controlling enzyme/substrate interaction, rather than directly influencing biochemical kinase activity.

